# The modular nature of long-association pathways in the human brain

**DOI:** 10.64898/2026.01.07.698021

**Authors:** Chiara Maffei, Marina R. Celestine, Ting Gong, Robert Jones, Evan Dann, Julia Lehman, Jean Augustinack, Hui Wang, Suzanne N. Haber, Anastasia Yendiki

## Abstract

Human brain functions are facilitated by large-scale networks, yet the organization of the long-range association pathways that form the wiring of these networks remains poorly understood. The advent of diffusion MRI (dMRI) tractography enabled, for the first time, the non-invasive mapping of these pathways, albeit at low resolutions. This contributed to a simplistic view of these pathways in the neuroimaging field, where they are conceptualized as monolithic entities, each linking a pair of distant cortical regions. This contrasts with classical anatomic tracer studies showing that each of these pathways is a network in itself, serving as a conduit for multiple fiber systems including short- and long-range projections. While higher-resolution dMRI and optical microscopy techniques for imaging the wiring of the human brain have emerged in recent years, our simplistic view of long association pathways from low-resolution tractography has not yet evolved. Here we present the first proof-of-principle study showing that, with these techniques, we can now replicate the modular nature of human brain pathways, as it has been known from tracer studies in animals. We demonstrate this on the most dorsal branch of the superior longitudinal fasciculus, the proposed backbone of the dorsal attention network. We show that the long-range, frontoparietal connections that we see with dMRI tractography at lower resolution (1.5 mm) are in fact composed of multiple, layered fiber systems of varying lengths that can be reconstructed with dMRI tractography at higher resolution (<750 μm). We validate this result across species, modalities, and scales by combining *in vivo* and *ex vivo* diffusion MRI with anatomic tracing in macaque and polarization-sensitive optical coherence tomography in human. These findings underline that advancing from macroscopic to mesoscopic imaging has the potential to transform our understanding of the wiring of the human brain, and allow the study of human brain networks in health and disease at a level of detail heretofore only possible in animal models.

## Introduction

Large-scale brain networks provide a fundamental framework for understanding human cognition and behavior. An accurate definition of the structural connections underlying these networks is essential for probing their organization in healthy individuals and detecting alterations associated with disease. The advent of diffusion MRI (dMRI) tractography allowed, for the first time, these structural connections to be characterized *in vivo* and non-invasively in the human brain. This set off extensive efforts to catalogue the main pathways of the brain and to come up with protocols for reconstructing them with dMRI tractography ^1,2^. In these efforts, perhaps due to the limitations of early dMRI acquisition and analysis methods, these pathways were conceptualized as *monolithic* structures, each connecting a pair of remote brain regions. This simplistic view of brain pathways prevails in human neuroimaging to this day.

In contrast, anatomy studies using tracer injections in non-human primates (NHP) show that each of these large white matter pathways functions as a conduit for a complex system of connections between multiple pairs of regions ^3,4^. Anatomic tracing allows us to follow axonal projections from their origin to their destination with microscopic precision. The resolution of conventional *in vivo* dMRI (1.5-3 mm), which is several orders of magnitude coarser than actual axons (1-10 μm), is insufficient for resolving this organization of white matter pathways in its full complexity. However, recent advances in dMRI ^5^ and emerging optical imaging techniques ^6^ are now making it possible to probe human brain circuitry at the meso- and micro-scale ^7^. Thus, it is worth revisiting the organization of human brain pathways, and examining whether these novel technologies are bringing us closer to the complex organization revealed by NHP tracer studies. Translating this finer-grained model of connectional neuroanatomy to the human brain is crucial, both for updating our understanding of the substrates of large-scale functional brain networks, and for guiding our *in vivo* analyses of these networks in patients.

In this work we demonstrate the feasibility of such a translation. We do this by combining: i) *Ex vivo* dMRI and tracer data in NHP (Macaque); ii) Multiscale *in vivo* and *ex vivo* dMRI data in humans; iii) Polarization sensitive optical coherence tomography (PS-OCT) in post-mortem human tissue. We show that dMRI tractography at mesoscopic resolution can resolve the modular nature of a long association pathway in the human brain that is lost at the lower resolutions of typical *in vivo* scans. We validate this modular structure by comparing the tractographic reconstructions to microscopic ground-truth tracer data in NHP and PS-OCT in human tissue.

This proof-of-principle demonstration focuses on the dorsal branch of the superior longitudinal fasciculus (SLF-I), a major long-range association fiber bundle broadly defined as connecting posterior parietal and dorsomedial frontal regions^8^. This pathway forms the structural backbone of the dorsal attention network (DAN), a well-known large-scale network that plays a key role in voluntary attention, spatial orientation, and action planning, while also encoding and maintaining preparatory signals and modulating top-down sensory regions^9^. Anatomic tracing studies in NHP describe the SLF-I as a system of fibers connecting multiple medio-parietal regions (areas PGm, PE, PEc, and 31) to different prefrontal regions (areas 6D, 8B, 9M, 9/46d)^8,10,11^. In humans, the morphology of the SLF-I remains controversial. Tractography and post-mortem dissection studies have yielded conflicting results; while some support direct, long, fronto-parietal connections, others support shorter projections or fail to identify the SLF-I as a distinct bundle^12–17^. These discrepancies between dMRI tractography in human and anatomic studies in NHP make the SLF-I an ideal testbed for this study.

Our results show that dMRI tractography of the SLF-I is dominated by long-range, frontoparietal connections at lower resolution (1.5 mm), but reveals a much more nuanced picture, with multiple fiber systems of different lengths, at higher resolutions (< 750 μm). These systems follow organizational principles that are homologous across NHP and humans: shorter-range connections that come in and out of the cortex rostral to area 6, and longer connections organized in layers between 6 and parietal regions. The neighboring cingulum bundle (CB), as opposed to the SLF-I, is shown to support most of the direct parieto-frontal connections. Anatomic tracing in NHP and PS-OCT in human tissue confirm this organization. These results demonstrate that imaging the wiring of the human brain at the mesoscale has the potential to reveal organizational principles that have been heretofore accessible only in animal models, as they cannot be resolved at conventional human neuroimaging resolutions. Importantly, this study provides a blueprint for revisiting other brain pathways in the future. A more refined understanding of human brain pathways as modular structures has the potential to reconcile structural and functional findings, and aid the interpretation of microstructural differences observed along these pathways in psychiatric and neurologic conditions.

## Results

### Integration of multi-scale connectional data across species and modalities

Figure 1 provides an overview of the different human and NHP datasets used in this work (see also Table 1 in methods section). The NHP datasets include: Anatomic tracing data from 7 animals (Dataset M1); *ex vivo* dMRI from 2 animals acquired at 700 μm (Dataset M2); *ex vivo* dMRI from 2 animals acquired at 500 μm (Dataset M3). The human datasets include: *Ex vivo* dMRI from two fixed brain hemispheres acquired at 750 μm (Dataset H1); *ex vivo* dMRI from 8 tissue blocks excised from the hemispheres in H1 and imaged at 250-300 μm (Dataset H2); polarization-sensitive optical coherence tomography (PS-OCT) from 10 small blocks further excised from the samples in H2 and imaged with an in plane resolution range of 3-10 μm (Dataset H3); *in vivo* dMRI from 1 subject acquired at 760 μm using an advanced multi-slab gSlider-SMS sequence^18^ (Dataset H4); *in vivo* dMRI from 10 subjects from the MGH-USC HCP acquired at 1.5 mm^19^ (Dataset H5). To ensure homology of dorso-medial brain regions, we adopted a common cortical parcellation and white matter dissection scheme across all datasets. Cortical regions were defined by combining manual labels from common cytoarchitectonic macaque atlases^20,21^ and automated parcellations in humans^22^ (Supplementary Table 1). To ensure precise assignment of connections, the white matter anatomical boundaries of the SLF-I and adjacent CB were outlined across datasets and modalities following the definitions and visualizations in^8^ (Figure 1).

**Figure 1.**
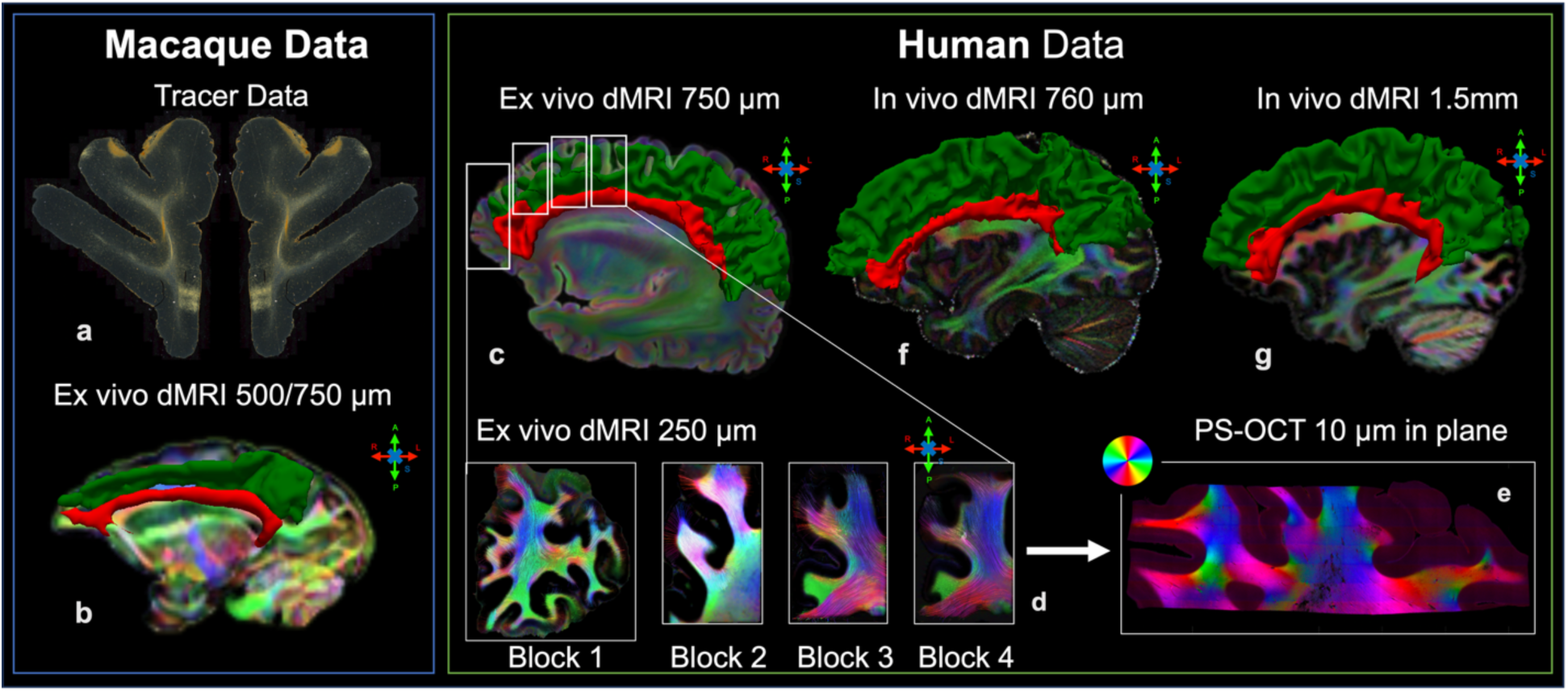
Dataset overview. The figure shows a representative image for each of the NHP (left panel), and human (right panel) datasets used in the study. The white matter boundaries of the SLF-I (green) and CB (red) are shown as 3D meshes for *ex vivo* NHP (**b**, Dataset M2), *ex vivo* human (**c-d**, datasets H1-H2), and *in vivo* human (**f-g**, datasets H4-H5) data, overlaid on directionally encoded color maps of the dominant orientation in the fiber orientation distribution functions (fODFs) from diffusion MRI. Color reflects the local orientation: red for left–right (mediolateral), green for anterior– posterior, and blue for superior–inferior, as indicated by the arrows next to the images. The approximate location of the four blocks (dataset H2) within the hemisphere (dataset H1) is shown in **c**. A representative photomicrograph from anatomic tracer data in macaque tissue (**a**) and an in-plane optic axis orientation map obtained with polarization-sensitive optical coherence tomography (PS-OCT) from one of the human tissue blocks (**e**) are also shown. In **e** colors represent axonal orientation within the imaging plane, as shown in the adjacent color wheel.

**Table 1.** Summary of datasets used in the study. The table lists the NHP and human datasets used in the study, along with their spatial resolution and the ID used to refer to each dataset throughout the manuscript. (dMRI: diffusion MRI; PS-OCT: polarization-sensitive optical coherence tomography; NHP: non-human primate).

| *NHP Data* |  |  |  | *Dataset #* |
| --- | --- | --- | --- | --- |
| **Anatomic tracer** | *Ex vivo* | 7 Cases |  | **M1** |
| **dMRI** | *Ex vivo* | 2 Samples | 700 $\mu\text{m}$ | **M2** |
| | *Ex vivo* | 2 Samples | 500 $\mu\text{m}$ | **M3** |
| **Human Data** |  |  |  | **Dataset #** |
| **dMRI** | *Ex vivo* | 2 Hemispheres | 750 $\mu\text{m}$ | **H1** |
| | | 8 Blocks | 250-300 $\mu\text{m}$ | **H2** |
| | *In vivo* | 1 Subject | 760 $\mu\text{m}$ | **H4** |
|  |  | 10 Subjects | 1.5 mm | **H5** |
| **PS-OCT** | *Ex vivo* | 10 Blocks | 3-10 $\mu\text{m}$ | **H3** |

### A fundamentally different organization in mesoscale versus macroscale tractography

We investigated the organizational principles of the SLF-I as revealed by tractography at different scales, by computing connectivity matrices between dorso-medial fronto-parietal cortical regions (Supplementary Table 1) from each whole-brain or hemisphere dMRI dataset included in the study (M2: 700 µm, M3: 500 µm, H1: 750 µm, H4: 760 µm, H5: 1.5 mm). For visualization, we generated connectograms (Figure 2), where the width of the lines is weighted by the number of streamlines linking each pair of cortical regions, and their color indicates the cortical areas they link, arranged in anterior-to-posterior order. For each dataset, we compared the connectogram including streamlines coursing through the SLF-I only, to the one including those coursing through both SLF-I and CB. Results show that at lower, more conventional dMRI resolution (H5: 1.5 mm) the SLF-I connectogram includes long, direct connections between frontal and parietal regions that are not present at higher resolutions. These long, direct connections between frontal (area 10, mPFC, lPFC) and parietal regions resemble the same connectivity profile of streamlines running within the CB. but run just above the CB. In both NHP and human high-resolution datasets (< 760 µm), most of the longer, direct connections between frontal (area 10) and pre-frontal cortex (PFC; Areas 9M, 8B) and parietal regions course within the CB and not the SLF-I, in accordance with the literature^23^. In NHPs the longest CB connections are between 9M and the parietal regions, whereas connectograms from human data also show connections with area 10. The SLF-I comprises shorter connections between adjacent regions, between pre-frontal and cingulate regions, between pre-frontal and pre-motor regions, and between pre-motor/motor and parietal regions. Of these, the highest number of streamlines is found for connections between pre-motor/motor and parietal regions. Figure 2 shows individual results for one representative subject from dataset H5, but the same profile is visible across the majority of the subjects, with the exception of one subject (Supplementary Figure 1).

**Figure 2.**
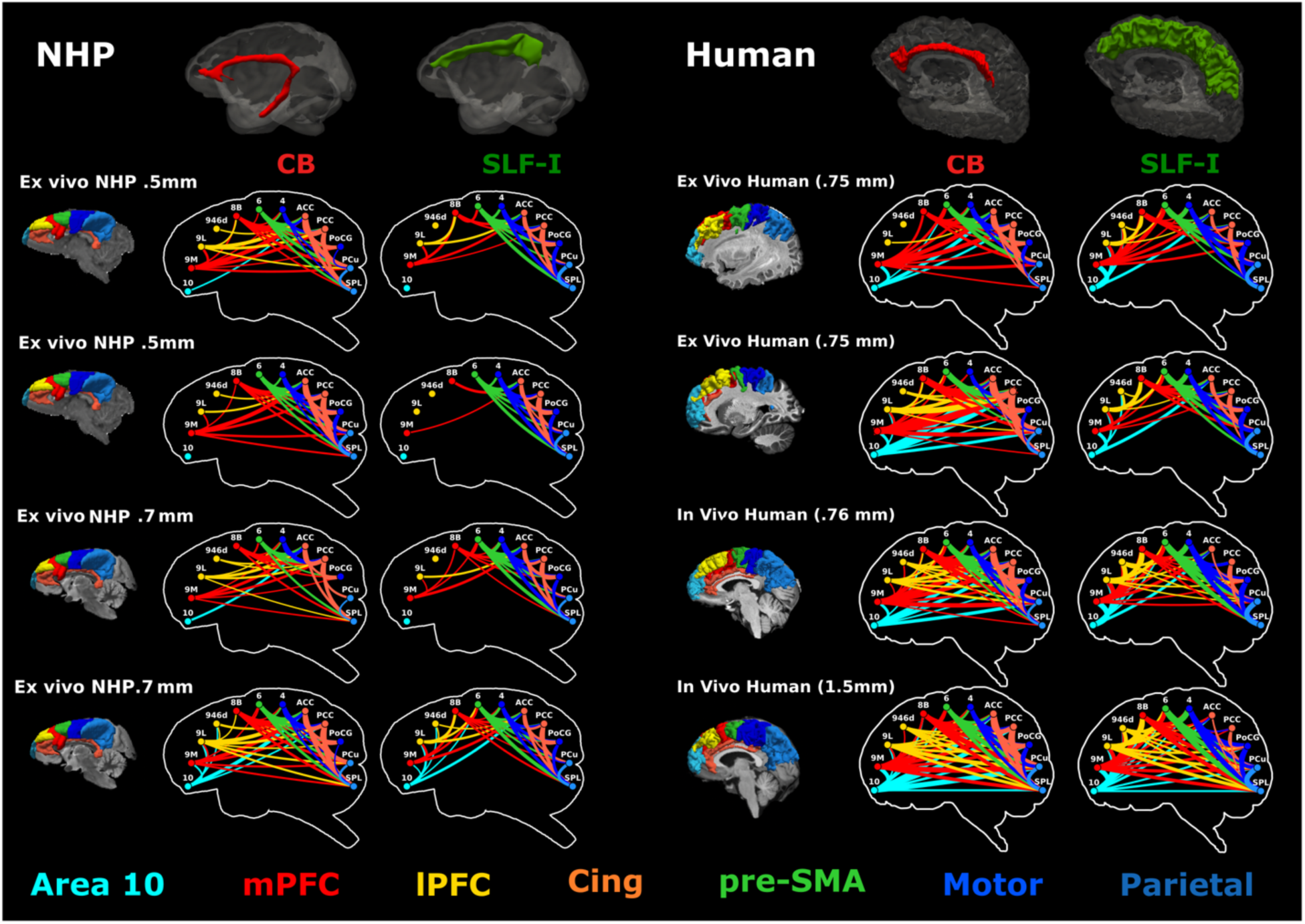
Connectivity of the dorsomedial cortex across scales and species. **Left)** Individual connectograms are shown for each of the four NHP datasets (M2, M3) for streamlines coursing through the CB (red) SLF-I (green). These inclusion ROIs are shown on top as 3D meshes for one of the datasets from M2. **Right**) Individual connectograms are shown for two hemispheres from dataset H1, for the whole-brain dataset from H4, and for one representative subject from dataset H5. The CB (dark red) and SLF-I (dark green) inclusion ROIs are shown on top as 3D meshes for one of the hemispheres from H1. The width of the lines is weighted by the number of streamlines between regions, and connection colors correspond to the cortical regions they link, arranged in anterior-to-posterior order: medial prefrontal cortex (mPFC, red) includes regions 9M and 8B; lateral PFC (lPFC, yellow) includes regions 9L and 9/46d; pre-supplementary motor area (pre-SMA, light green) corresponds to region 6; motor areas (blue) include region 4 and regions 1,2,3 in the posterior central gyrus (PCG) ; cingulate regions (orange) include anterior (ACC) and posterior cingulate cortex (PCC) (Areas 24, 32, 8/32, 9/32, 6/32, 23, 31); parietal region (purple) include the superior parietal lobule (SPL) and the precuneus (PCu). For each dataset, the parcellation of the dorsomedial cortical regions is shown on the left using the same color scheme as the connectograms.

### High-resolution *ex vivo* dMRI reveals the modular organization of the SLF-I in humans

To investigate the mesoscopic organization of the SLF-I in greater anatomical detail, we performed manual virtual tractography dissections of the SLF-I from both high-resolution *ex vivo* dMRI and PS-OCT in two human hemispheres (H1, 750 μm; H2, 250 μm; H3, 10 µm).

Results from H1 (750 μm) show that the frontal portion of the SLF-I (Rostral to Area 6) is largely composed of short relay fibers that enter and exit the white matter territory of the SLF-I to connect different dorsomedial pre-frontal cortical regions (Figure 3). For both samples, we observed longer direct connections between 6M and the SPL and precuneus (that were more extensive in the latter). These streamlines are organized in progressively shorter bundles that layer on top of each other (Figure 3). Rostral to area 6M we observed a majority of shorter connections between 6M and 8B, 8B and 9M, within area 9M, and between area 9M and 10, with very similar anatomical profiles between the two samples. Only a few longer connections between parietal regions and 8B or caudal 9M were observed to run underneath these shorter connections and above the CB.

**Figure 3.**
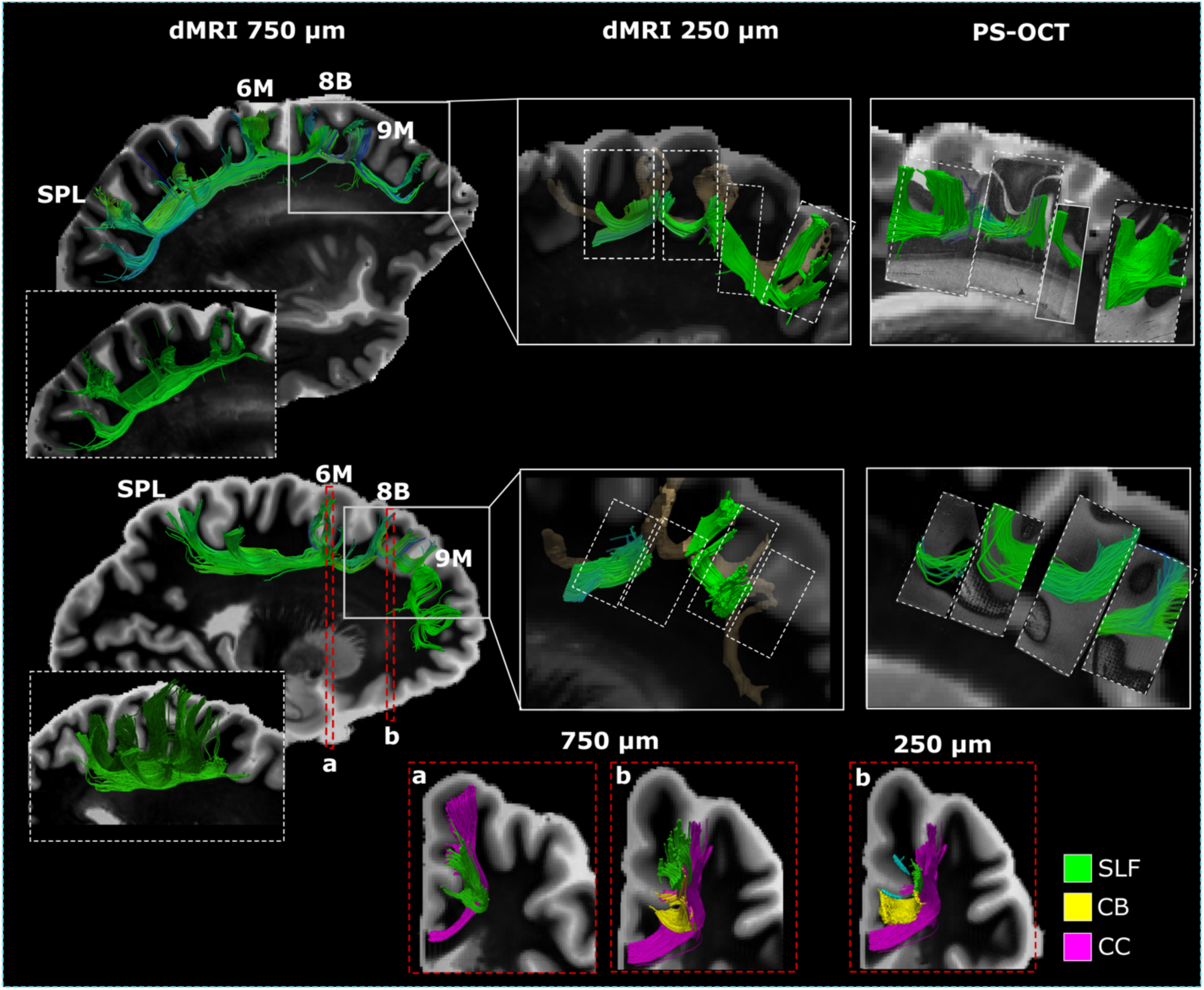
Diffusion MRI tractography in humans. Manual virtual dissections of the SLF-I are shown in sagittal view for two hemispheres, for dMRI data of the whole hemisphere acquired at 750 µm (Dataset H1), dMRI data of tissue blocks acquired at 250 µm (Dataset H2), and PS-OCT data of tissue blocks acquired at 10 µm in plane (Dataset H3). For each hemisphere, dashed white boxes show the layer organization of the SPL-6M streamlines. Solid white boxes indicate the approximate location of the tissue blocks from H1 used in datasets H2 and H3. Tractography dissections from 250 µm dMRI data are shown overlaid on 3D volumes of the dissections obtained from the hemisphere data (in light brown). Dashed white boxes indicate the approximate locations of the PS-OCT blocks. At the bottom of the image, panels a and b show coronal views of the relationship between the SLF-I fibers and other white matter bundles for different locations (6M, 8B) and different resolutions (750 µm, 250 µm). CB: cingulum bundle; CC: corpus callosum; SPL: superior parietal lobule.

Tractography generated from higher resolution dMRI data (H2, 250 μm) and PS-OCT (H3, 10 µm) closely resembled the short connection profile observed at 750 µm for regions rostral to 6M, and no long direct connections were identified running underneath these fibers from either modality. Compared to more posterior areas (Figure 3a), at the level of area 8B, the SLF-I lies superior to a dense layer of streamlines leaving the SFG to enter the CB. These streamlines occupy a large portion of the white matter lateral and superior to the cingulate sulcus. Tractography at higher resolution (250 µm) allowed us to better distinguish the different fiber population within this white matter region showing a thin layer of streamlines running between the SLF-I and the cingulate sulcus and entering the CB (Figure 3, bottom).

### PS-OCT in human tissue reveals predominantly short-range connections rostral to area 6

Figure 4 shows en-face optic axis images in sagittal planes, illustrating the orientation of white matter fibers within the SFG across blocks. Multiple sagittal sections illustrate different fiber organizations from more medial (Figure 4a) to more lateral (Figure 4c). At the level of the cingulate sulcus (CS) (Figure 4a), CB fibers are clearly visible running within the cingulate gyrus in all blocks. At this level, fibers within the SFG run superior the CS following the gyral anatomy within all blocks. In figure 4b, at the medial end of the CS and close to the medial wall of the SFG, fibers within the most posterior block (located at the level of rostral area 6 and area 8B) run along the anterior-posterior axis and either turn to project dorsally to the cortex or continue to enter the adjacent block (within 9M). In 9M most of the fibers project dorsally without further continuing anteriorly into rostral 9M. While no anterior-posterior fibers are visible superior to the CS, CB fibers are clearly visible continuing inferiorly within the supero-medial cingulate gyrus. As we move more lateral, within the white matter of the SFG, regions within 9L and 8B are dominated by fibers running inferior-superior (Figure 4c), including fibers leaving the CB and projecting dorsally (Figure 4c, central block). Contrary to tractography results obtained at 750 μm, en-face PS-OCTT data showed no longer connections from 8B or caudal 9M to parietal regions.

**Figure 4.**
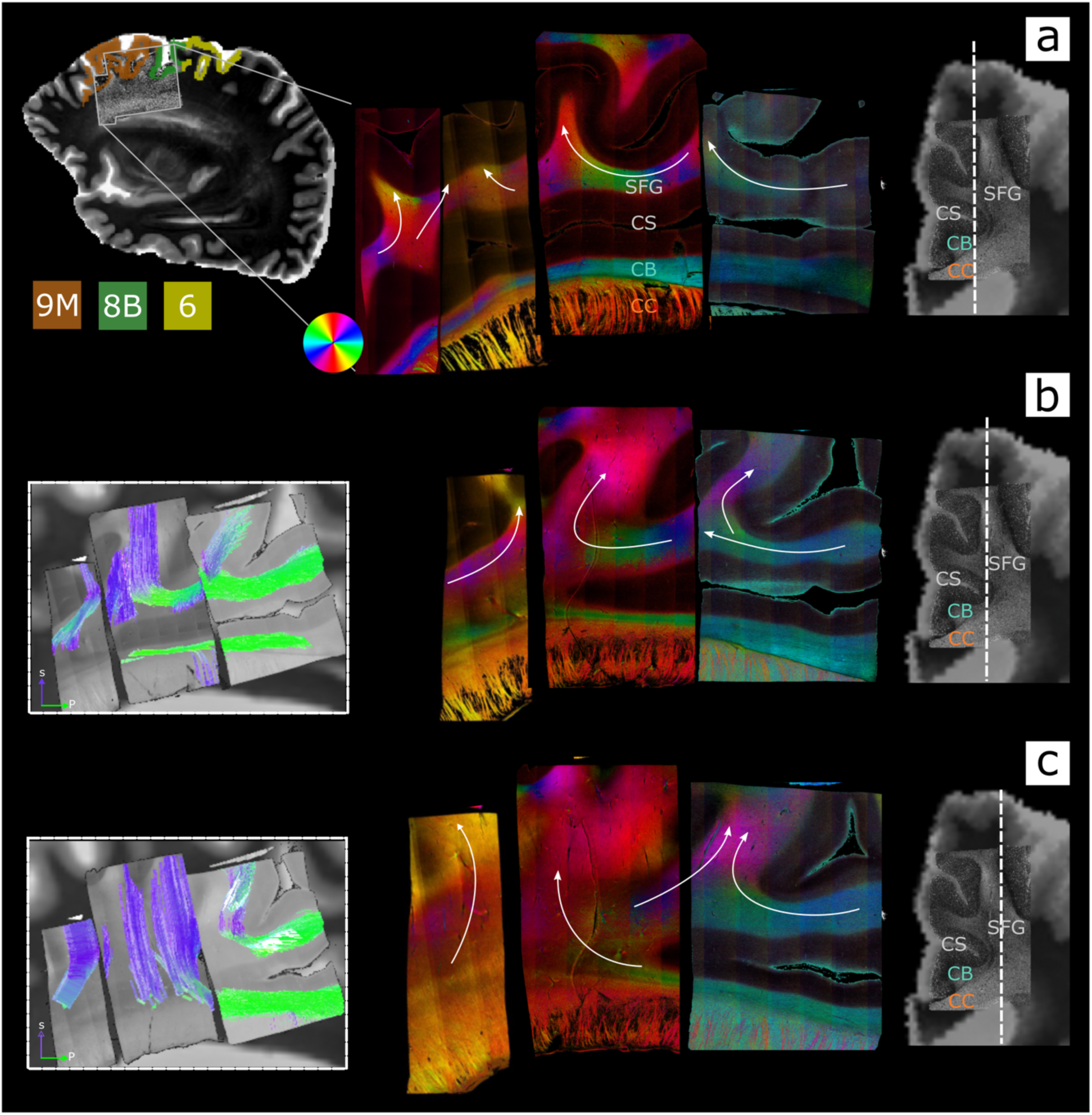
PS-OCT in humans. Sagittal en-face images of optic axis orientations in the superior frontal gyrus (SFG) from PS-OCT are shown for one human sample at three different levels: **a)** most medial, over the cingulate sulcus; **b)** more lateral, at the medial end of the cingulate sulcus; **c)** most lateral, within the SFG white matter. To the right of each row, coronal retardance images from PS-OCT are overlaid on the co-registered dMRI data and white dashed lines on coronal sections indicate the respective sagittal level. Labels for the superior frontal gyrus (SFG), the cingulate sulcus (CS), the cingulate bundle (CB), and the corpus callosum (CC) are added to each coronal image and on one sagittal PS-OCT panel (top). The location of the PS-OCT samples within the hemisphere is shown in the top left together with the outline of the cortical regions they include (9M, 8B, rostral 6). For each level, white arrows are overlaid on the PS-OCT images to indicate the overall orientation of the fibers. The optic axis orientation colormap is defined by the color wheel displayed next to **a** (top left). For levels **b** and **c** tractography performed on the PS-OCT orientations is shown cropped to the displayed slice. CB: cingulum bundle; CC: corpus callosum; CS: cingulate sulcus; SFG: superior frontal gyrus.

### Anatomic tracing in NHP shows multiple fiber systems of different length within the SLF-I

We charted the trajectory of the SLF-I fibers in 7 male macaque monkeys (Supplementary Table 2) with injections along the rostro-caudal axis of the dorso-medial cortex (see Methods, Figure 8). Axonal projections were classified as SLF-I fibers if they coursed anterior-posterior within the dorso-medial white matter boundaries previously described for the SLF-I^8^. For all 7 cases, fibers enter the SLF-I from injection sites on the medial (Cases 1-5) or lateral (Case 6 and 7) wall of the dorsomedial cortex and travel within the SLF-I before terminating in the dorsomedial (SLF-I terminals) or adjacent dorsolateral cortex. Some fibers travel within the SLF-I for a short distance before exiting it to enter other white matter bundles like the CB, corpus callosum (CC), or striatal white matter through the subcortical bundle (SB). Figure 5 shows the location of injection sites and SLF-I terminals for all cases, and the entire SLF-I fiber trajectories for three cases (Cases 1,6,7). The longest direct SLF-I connections were observed between parietal regions PE/PGM and pre-SMA regions (6M, 6D) (Figure 5; cases 6 and 7). Injection sites rostral to 6 (9M and 8B) showed only short-range fibers within the SLF-I terminating primarily in 9L, 6M, 6D, and 24 (Cases 1-3) with most fibers entering the CB (Figure 5a, 5b). Injections in rostral 6M showed only moderate sparse terminations in 9M and 6D with most of the fibers traveling towards subcortical and lateral regions (Cases 4 and 5). In Case 1 fibers leave the injection site in 9M and enter the SLF-I medially (Figure 5a); they travel posterior for a short distance within the SLF-I and terminate in 8B, 6D (Figure 5b), and 6M (Figure 5c). Along this course, some fibers also leave the SLF-I to terminate laterally in 9L or enter the CC (Figure 5b), CB or SB (Figure 5c). In Case 6 fibers leave the injection site in 6D to travel both anteriorly and posteriorly within the SLF-I; anteriorly, fibers terminate in 8B and 9M (Figure 5d); posteriorly, fibers terminate in PGM and PE (Figure 5g). Similar to Case 1, fibers leave the SLF-I to enter subcortical regions at the level of 6M (Figure 5e), and the CB at different levels (Figure 5f). In Case 7, the only case with an injection in the parietal lobe, fibers enter the SLF-I laterally in PE (Figure 5l) and travel anteriorly within the SLF-I until they terminate in caudal 6M (Figure 5h). Like in the other cases, fibers leave the SLF-I to enter the CB, CC or lateral regions but also show moderate and light terminal fields in sensory motor regions 3 and 4 (Figure 5i). In all cases, SLF-I fibers travel superior to the cingulate gyrus on the medial wall of the SFG. When present, fibers projecting to and from the CB are always found to be medial to the SLF I fibers, running in a thin layer very close to the cortex. In support of the tractography results, in 9M most of the fibers project dorsally without further continuing anteriorly into rostral 9M. In none of the cases did we observe fibers going directly connecting PGM/PE to area 8B or 9M, *i.e.*, the longer-range frontoparietal connections often found in SLF-I definitions in the dMRI literature and that we observed at lower *in vivo* dMRI resolutions (Figure 2).

**Figure 5.**
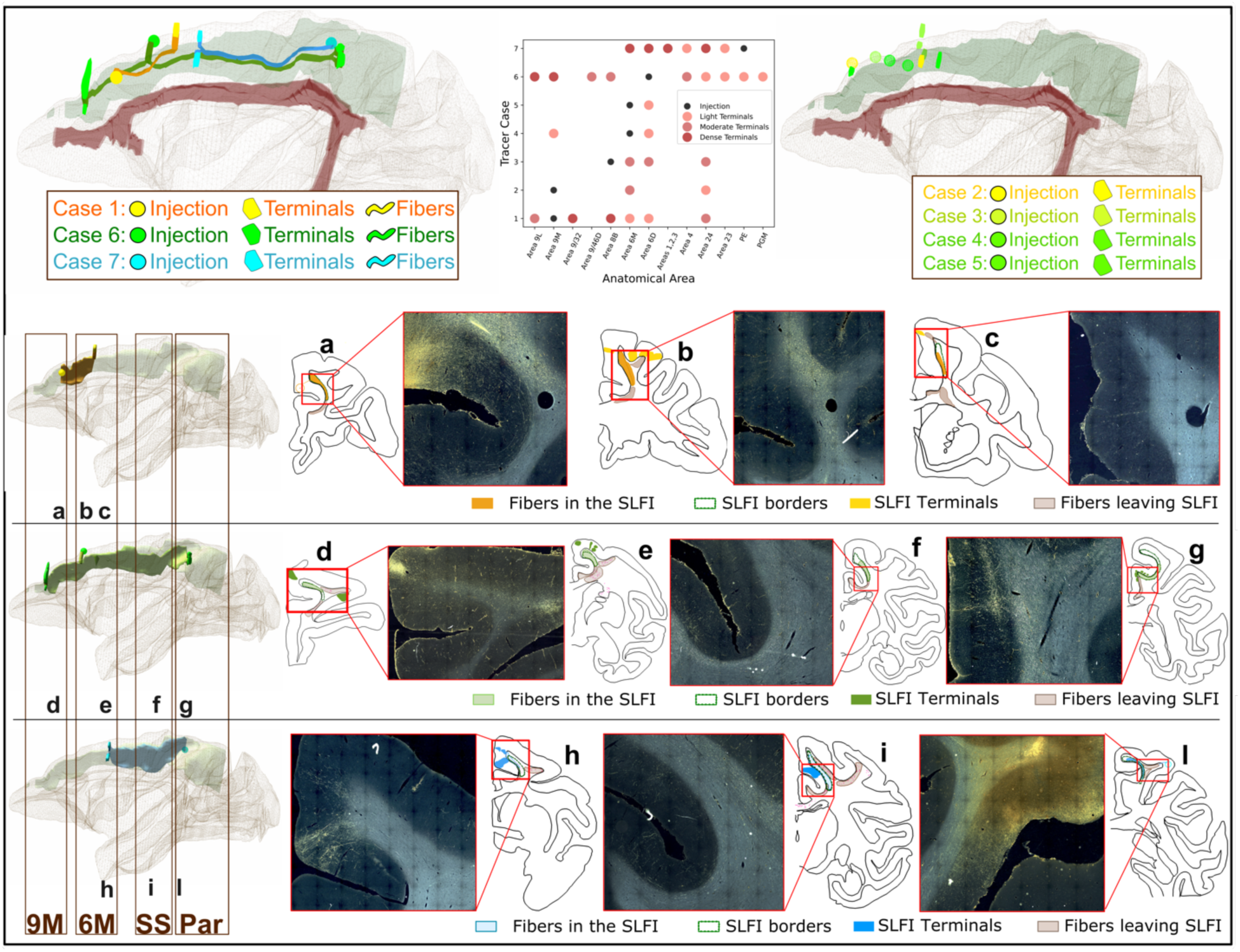
Anatomic tracer injections in NHP. **Top left:** 3D model of fiber pathways running within the SLF-I in common template space from the 3 fully charted cases (Case 1: orange, Case 6: green, Case 7: blue). The injection sites are shown as spheres, along with the centroids of the fiber trajectories and the fiber terminals located most distant from the injection site. For reference, 3D reconstructions of the SLF-I and CB bundles, based on the anatomical definition from ^8^, are also shown in green and red, respectively. **Top right**: A 3D model of injection sites and terminal fields in template space is shown for the four remaining cases (2-5). **Top center:** A scatter plot shows the injection site and location of terminal fields (light, moderate, and dense) for all 7 cases. **Bottom**: 3D models of the full extent of fiber trajectories in individual space are shown (left column) for each of the 3 fully charted cases along with 2D outlines (**panels a-l**) of representative slices within 9M, 6M, somatosensory (SS), and parietal (Par) regions showing the fibers traveling within the SLF-I, fibers leaving the SLF-I to enter other bundles, and SLF-I fiber terminals. The borders of the anatomical definition of the SLF-I ^8^ are also shown (green outlines). For most slices the corresponding photomicrographs are also shown.

### *Ex vivo* tractography in NHPs replicates the modular organization of SLF-I connectivity

Manual virtual dissections of the SLF-I using dMRI tractography data in two *ex vivo* macaque brains (one from M2 (700 µm); one from M3 (500 µm)) show streamlines within the SLF-I with similar connectivity patterns to the tracer data (Figure 5a) and the *ex vivo* tractography in humans (Figure 3). Most of the reconstructed streamlines run between PE/PGM and area 6M. Rostral to area 6M, the SLF-I is comprised mostly of short streamlines connecting 6M and 8B/9M. In contrast to the tracer data, tractography shows some longer streamlines connecting PGM/PE and area 8B (M2, M3) and to a lesser extent area 9M (M2) (Figure 6a). These streamlines run inferior to the shorter connections between 6M and 8B/9M. In both cases, the streamlines running between PGM/PE and 6M are organized in layers of progressively shorter connections (Figure 6a), like in the human *ex vivo* tractography data (Figure 3). In agreement with the tracer data, at the level of 6M we could delineate streamlines leaving the SLF-I to enter the CB, CC, or to project to lateral cortical regions (Figure 6b). In contrast to the tractography results in humans, the SLF-I streamlines run in a more superior location within the SFG. To compare the connectivity patterns of the SLF-I between the tracer and tractography data, we placed ROIs in the same anatomical location as the injection sites for Cases 1, 6, and 7, and isolated streamlines running within the SLF-I. Figure 6c shows that tractography replicates the connectivity profiles observed in the tracer data: for Case 1 (yellow), streamlines run from 9M to 6M; for Case 6 (green), streamlines run from 6D to 9M and from 6D to PE; for Case 7 (blue), streamlines run from PE to caudal 6M. To evaluate the overall distribution of connections from the injection point, we compared individual tracer results to tractography seeded at the same anatomical location as the injection site in Cases 1 (9M), 2 (9M), 6 (6D), and 7 (PE) (Figure 6d). For seed ROIs in 9M (Cases 1, 2), streamlines have very similar connectivity patterns projecting heavily to lateral (9L, 9/46) and subcortical areas, and CB directly from the injection region. In agreement with tracer experiments, few streamlines enter the SLF-I from these cortical areas and run only a short distance within it before terminating in 6M. In contrast to the anatomic tracer, where axons traveling within the SLF-I occupy the entire extent of the medial white matter superior to the cingulate sulcus, SLF-I streamlines occupy the most superior part of the SFG (green streamlines in Figure 6d; y=29, 33). The white matter inferior to the SLF-I streamlines and superior to the cingulate sulcus is instead dominated by streamlines running inferior-superior, mainly projecting to the CC and CB.

**Figure 6.**
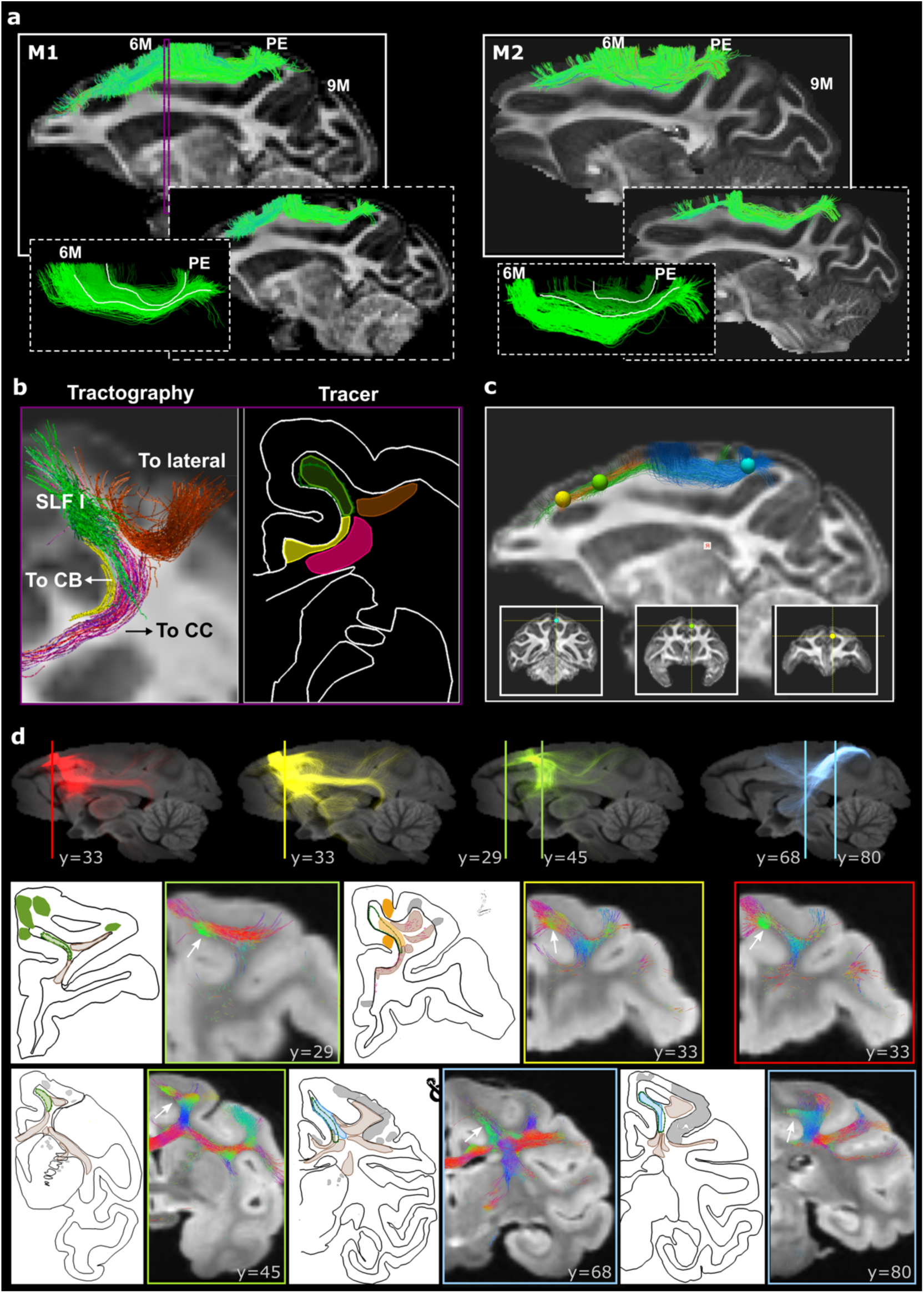
Diffusion MRI tractography in NHP. **a)** Manual virtual dissections of the SLF-I for two macaques (M2, 700 µm; M2, 500 µm). For each macaque, dashed white boxes show the subcomponents of the SLF-I connecting PE with 6M and 6M with 8B (right), and the layer organization of the PE-6M streamlines (left). Tractography is shown superimposed to FA maps. **b)** The relative position of the SLF-I with adjacent white matter bundles (CC, pink; CB, yellow; lateral projections, orange) is shown for a coronal section at the level of 6M for dMRI tractography (M2) and tracer data (Case 6). The location of the dMRI slice is indicated by the purple rectangle in A. **c)** SLF-I tractography streamlines identified by spherical ROIs placed at the same anatomical location as the injection site for Cases 1 (yellow), 6 (green), and 7 (blue). The exact location of the spherical ROIs is shown underneath in coronal view. **d)** All the streamlines obtained when seeding tractography from spherical ROIs placed at the same anatomical location as the injection site for Cases 1 (Yellow), 2 (Red), 6 (Green), and 7 (Blue). Tractography results are shown for different coronal slices (indicated by the Y value) and different seed ROIs (indicated by the color of the box outline), and compared to individual tracer injections for the corresponding Case at a similar anatomical location. The location of the coronal slices is indicated by the vertical line shown in sagittal view. Tractography streamlines are color-coded based on their orientation: green for anterior-posterior, red for right-left, blue for inferior-superior. White arrows indicate green streamlines running anterior-posterior within the SLF-I. For the tracer charts, the same colors are used as in Figure 2, with SLF-I fibers and SLF-I terminals colored based on Case (Case 1: Yellow; Case 6: Green; Case 7: blue), fibers leaving the SLF-I in brown, and all other terminals in grey. CB: cingulum bundle; CC: corpus callosum.

Tractography results also consistently show streamlines projecting laterally to the middle and inferior frontal gyri that are not visible in the tracer experiments (Figure 6d; y=33, 45). For ROIs placed in 6D and PE (Cases 6, 7), tractography results are very similar to the tracer data. SLF-I streamlines occupy the medial white matter superior to the cingulate sulcus and streamlines leave the SLF-I to enter the CC, CB, and subcortical fibers (Figure 6d; y=45, 68, 80).

### High-resolution tractography in NHPs and humans replicates connectivity from tracer data

To compare tractography in both NHPs and humans with ground-truth tracer results, we binarized the connectograms by aggregating results from all 7 tracer cases and isolated those streamlines in each tractogram that connected the regions in which we have tracer injections: PFC, pre-SMA, and parietal regions (Figure 7). As expected from the NHP virtual dissections (Figure 5d), streamlines from PFC (9M and 8B) that run within the SLF-I mainly project to adjacent regions anterior or within the pre-SMA, in agreement with the tracer data. Notably, *ex vivo* macaque tractography fails to reconstruct streamlines between areas 9M and 6, whereas human tractography shows false positive connections to cingulate and motor cortex at 750 µm, and also to parietal regions at 1.5 mm. Streamlines from pre-SMA correctly project to both anterior and posterior regions in NHP and human tractography; however, high-resolution *ex vivo* tractography misses lPFC projections and falsely identifies PoCG connections absent in tracer, while *in vivo* tractography interestingly misses connections to the cingulate areas while adding connections to area 10. Connectograms from parietal regions are identical between tracer data and ex vivo tractography in NHP, while they show false negative connections to anterior cingulate regions for ex vivo and a large number of false positive connections to frontal regions in vivo for human data (Figure 7). For parietal regions, tracer and *ex vivo* tractography results were consistent in NHP, whereas human data showed false negative connections to anterior cingulate cortex ex vivo and numerous false positive frontal connections in vivo (Figure 7).

**Figure 7.**
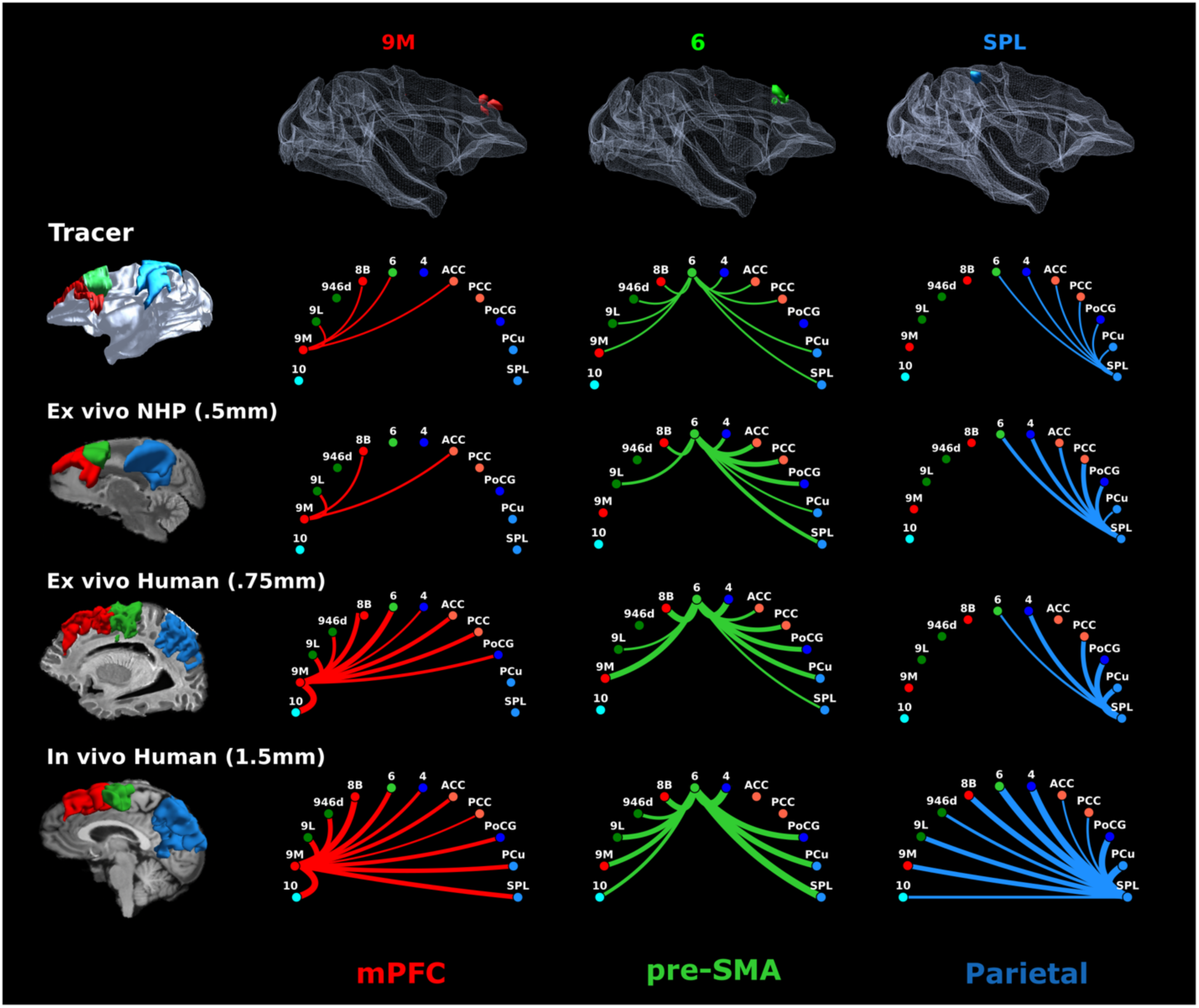
Connectograms of regions 9M, 6, and SPL across modalities, species, and scales. Connectograms for the tracer data, the NHP dMRI (M3), and the human dMRI (H1, H5) are shown for the three cortical regions for which we have tracer injections: area 9M (red), area 6 (green), SPL (blue). Connectograms from the tracer data are binary (line indicates fiber connections between two regions in at least one of the cases) and were created by aggregating results from the 7 tracer cases. Injection locations for all cases are shown in 3D on top overlaid on a white matter mesh in template space. Cortical regions used to assign terminations are shown on the left. For dMRI data these regions were also used as seeds. The width of the lines for dMRI connectograms is weighted by the number of streamlines between regions, and connection colors correspond to the cortical regions they link, arranged in anterior-to-posterior order: medial prefrontal cortex (mPFC, red) includes regions 9M and 8B; lateral PFC includes regions 9L and 9/46d; pre-supplementary motor area (pre-SMA, light green) corresponds to region 6; motor areas include region 4 and regions 1,2,3 in the posterior central gyrus (PCG) ; cingulate regions include anterior (ACC) and posterior cingulate cortex (PCC) (Areas 24, 32, 8/32, 9/32, 6/32, 23, 31); parietal region include the superior parietal lobule (SPL) and the precuneus (PCu).

## Discussion

Here we combine data across scales, modalities, and species to provide an updated anatomical representation of long association pathways of the human brain. By leveraging high-resolution *ex vivo* dMRI we bridge the gap between microscopic and macroscopic anatomical representations of the SF-I, paving the way for a more detailed and rigorous definition of other major white matter bundles. High-resolution dMRI (< 750 µm) reveals that the direct, long-range connections dominating SLF-I reconstructions at typical *in vivo* dMRI resolutions (1.5 mm) are composed of shorter fiber systems that closely resemble the organization observed in tracer data. The direct long connections between parietal and frontal regions course within the CB, rather than the SLF-I, in agreement with the literature^23^. Across modalities and species, we reveal a consistent modular organization of connections within the SLF-I, with short-range connections dominating the SLF-I rostral to area 6 (pre-SMA), and longer, direct connections between parietal regions PGM/PE and area 6, organized in layers of progressively shorter connections. Our results offer a refined anatomical definition of the SLF-I that reconciles older anatomical debates across modalities and species, and provide a revised framework for studying major association bundles of the human brain using tractography techniques.

### Limitations of conventional imaging and dissection techniques

High-resolution dMRI tractography shows the SLF-I as a system made of multiple fiber populations of different lengths, challenging the simplistic view of white-matter pathways that is prevalent in human neuroimaging. This complexity, that is independently supported by anatomic tracer data in macaques and PS-OCT in humans, is missed by *in vivo* lower-resolution dMRI tractography, which generates several artifactual, direct fronto-parietal connections. This observation helps explain the discordant results of previous tractography studies in humans, which were confounded by low dMRI resolution. While those studies overall agreed on the posterior terminations of the SLF-I in the SPL and precuneus, the exact location of its anterior terminations has been a matter of debate. Some tractography studies have showed them extending anteriorly to connect regions in the dorsal prefrontal (areas 8B, 9, 46) and anterior cingulate cortex (area 24), while others showed them constrained to the rostro-medial part of area 6 (SMA and pre-SMA). Because of its proximity to the CB, some tractography studies even questioned the existence of the SLF-I as a distinct fiber bundle and suggested it should instead be considered part of the cingulum fiber system ^15,24^. Gross dissections in humans report similar discordant results, some classifying the SLF-I as a frontoparietal long association bundle ^25^, some suggesting that frontal and parietal regions are connected instead by a succession of short association pathways^16^, and some more recent evidence describing two separate but contiguous SLF-I segments joining in the rostral part of area 6^26^.

These contradicting results on the morphology of this pathway among studies using *in vivo* tractography or *ex vivo* blunt dissections reflect limitations of both techniques in correctly disentangling different white matter fibers in complex regions. At lower resolutions (including the resolution of Klinger’s gross dissections^16^) the spatial vicinity of SLF-I and CB may give the erroneous impression of the existence of a long association pathway in both tractography and post-mortem dissection studies^16^. Over some distances, the shorter components of the SLF-I contain fiber populations with very similar orientations to the CB, creating challenging “bottleneck” configurations that at low tractography resolutions foster error propagation ^27^ and yield consistent false positive reconstructions^28^.^26^

### Mesoscopic imaging reveals the modular nature of human white-matter pathways

High-resolution tractography resolves these distinct fiber populations, showing the SLF-I as a modular system composed of shorter-range connections of different length that run beneath the superficial white matter and separate from the CB trajectory to connect different regions within the prefrontal, premotor, motor, and parietal cortices. This organization becomes increasingly apparent at 250 μm (Figure 3) and is independently supported by PS-OCT measurements (Figure 4). By integrating multiscale cross-species data, we capture the true complexity of the SLF-I, highlighting the limitations of conventional dMRI and reconciling prior conflicting definitions by revealing that they were in fact referring to subcomponents of a common, more complex system.

### Common organizational principles across species

Combining data on connectional anatomy across species allowed us to compare the fiber systems comprising the SLF-I and identify conserved organizational principles between NHP and human brains. In both species, fibers leaving cortical regions rostral to the pre-SMA mainly project to lateral (areas 9/46, 9L), cingulate (areas 24, 32, 23) and subcortical regions, with fibers travelling only a short distance within the SLF-I boundaries (Figures 2,4). While tractography shows a few streamlines connecting parietal and frontal regions (areas 8B, 9M) in both humans and NHPs, most of the longer, direct streamlines course between parietal regions and area 6. In both species, *ex vivo* tractography shows that these longer connections are organized in layers of progressively shorter connections (Figures 3, 5). Compared to humans, tracer data and *ex vivo* tractography in NHPs show SLF-I fibers running within a more dorsal location within the SFG, possibly reflecting increased cortical gyrification in humans that displaces homologous fibers inferiorly toward the deep medial frontal white matter. While the location of SLF-I fibers within the superior frontal gyrus differs across species, the overall organizational commonalities highlight the importance of tracer injection studies in NHPs for validating dMRI and exploring evolutionarily conserved structural–functional principles within the dorsal attention network^29^.

### A refined anatomical framework for connectional anatomy in the human brain

The multiscale approach adopted here on the SLF-I establishes an updated anatomical framework for investigating the internal organization of major white matter pathways in the human brain. Our results support a modular organization in which large bundles are composed of different fiber systems of different lengths. This organizational principle is consistent with observations from prior tracer studies in NHPs, which have shown that axons use major association bundles as conduits, traveling within them for part of their trajectory before diverging toward their cortical targets^3,4^. This general organization gives rise to several branching configurations, where fibers consistently fan away from major bundle trajectories. Direct comparison of tracer and tractography data in NHPs has previously shown that these configurations are a common source of errors for dMRI tractography^30^. The present findings extend this principle to the human brain and demonstrate that its representation from dMRI tractography is dependent on spatial resolution. At lower, conventional resolutions (1.5 mm) the reconstruction of the SLF-I results mainly composed of long-range connections, resembling the same organization of the adjacent CB. At 750 μm, tractography begins to reveal the internal organization of SLF-I, resulting in a significant reduction of the long-to-short connection ratio relative to the 1.5-mm reconstruction. These results are consistent with the broader anatomical observation that much of cortical connectivity is relatively short-range^31,32^.

A refined anatomical framework of major white matter pathways is essential a better understanding of how the wiring of the brain gives rise to large-scale functional networks^33^. In contrast to the conventional view of white matter pathways as monolithic structures, each of them is comprised of subcomponents that can participate in multiple such functional networks. In the case of the SLF-I, this more detailed characterization clarifies how different fiber populations may support both dorsal attention and frontoparietal control networks, and how specific subcomponents may become selectively vulnerable in disease. Understanding the modular organization of major pathways is also important for interpreting localized microstructural differences that are often observed at different sections along major association pathways in both psychiatric and neurologic conditions^34^.

This study illustrates the potential of multiscale, multimodal imaging for revolutionizing our understanding of the connectional anatomy of the human brain. Our results suggest that spatial resolution influences not only the precision with which a known pathway is reconstructed, but also the anatomical model of that pathway that emerges from that reconstruction. From this perspective, the long trajectories conventionally used to define association bundles may represent only one level of their organization. Scaling up this approach to perform a systematic mapping of long-association pathways across the entire brain is necessary to determine the organizational principles of other major fiber systems beyond the SLF-I, and provide a reference for evaluating *in vivo* tractography. This goal aligns with ongoing large-scale efforts to map the full wiring diagram of the primate brain^35^. Such efforts will catalyze the introduction of models of white-matter pathways that explicitly account for multiscale organization. This is a necessary step towards a meaningful integration of microstructural, macroscale, and functional data to investigate how circuits are organized, and how this organization influences cognition, perception, emotion, and behavior.

## Methods

The different human and NHP datasets used in this work are listed in Table 1. All dMRI datasets underwent the same basic preprocessing steps: they were denoised and corrected for motion and eddy current-induced distortions. For each dMRI dataset, estimations of local fiber orientations used for tractography analyses were obtained by constrained spherical deconvolution (CSD)^36,37^. Cortical parcellations of the frontal and parietal cortex were obtained for each dataset to enable anatomical comparisons across modalities and species. Acquisition and processing details for each dataset are provided below.

### Anatomic Tracer Data in NHPs: acquisition and preprocessing

Figure 8 shows the location of the injections for the 7 cases used in this study (See also supplementary table 1). Cases were selected based on injection location, presence of fibers within the SLF-I, and overall quality of the transport. Injections with tracer leaking into adjacent cortical regions or into the white matter were excluded. Surgery and tissue preparation were performed at the University of Rochester Medical Center and details of these procedures were described previously ^3,30,38^. Briefly, each monkey received a 40–50 nl. injection of an anterograde/bidirectional tracer (lucifer yellow (LY), fluororuby (FR) or fluorescein (FS)) conjugated with dextran amine (10% in 0.1 M phosphate buffer, pH 7.4; Invitrogen). Tracers were pressure injected over 10 minutes using a 0.5 μl Hamilton syringe. Following the injections, the syringe remained in situ for 20 minutes to avoid leakage and contamination as the needle was withdrawn. Twelve to 14 days after the injection, animals were deeply anesthetized and perfused with saline followed by 4% paraformaldehyde/1.5% sucrose solution. The brains were removed, postfixed overnight and cryoprotected in increasing gradients of sucrose (10, 20, and 30%). Specimens were sectioned in 50 μm-thick coronal slices on a freezing microtome into 0.1_M_ phosphate buffer or cryoprotectant solution as previously described^39^. To visualize the tracers, one in every 8th section was processed for immunocytochemistry resulting in an inter-slice gap of 400 μm. Additional details on the surgical and histological procedures can be found in previous papers ^3,40^. All experiments were performed in accordance with the Institute of Laboratory Animal Resources Guide for the Care and Use of Laboratory Animals and approved by the University of Rochester Committee on Animal Resources.

**Figure 8.**
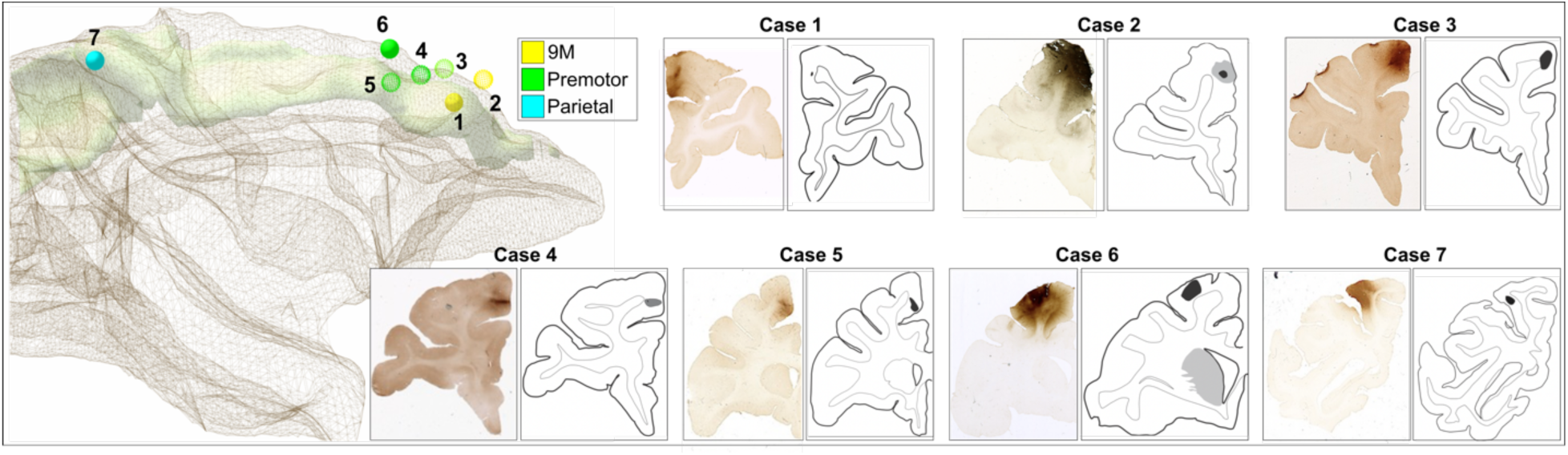
Tracer injection sites. Left: A 3D model of an in-house template shows the 3D white matter mesh (brown) overlaid with the location of the injection sites, along with the SLF I anatomical reconstruction (green) from ^8^. The center of the injection site is placed within the model and visualized as a sphere of equal radius across cases, with color indicating anatomical location (Yellow: 9M, green: premotor; blue: parietal). Cases for which we charted fibers serially on every slice are shown as filled spheres (Cases 1,6,7). **Right**: Micrographs and schematics of coronal sections show the injection location for each case. Case numbers correspond to Supplementary Table 2.

Sections were viewed under dark-field illumination and images were obtained using Neurolucida software (MBF Bioscience). Fiber bundles were outlined using a 4.0x or 6.4x objective. Some individual fibers within each bundle were charted at higher magnification (10x) to indicate directionality. We followed axonal projections as they left the injection site, travelled through the dorso-medial cortex, and reached their termination points within the frontal, premotor, motor, cingulate, or parietal regions. Fibers traveling together with discernible boundaries were outlined as a bundle. For both fibers bundles and terminal fields, the fiber density was assessed visually to classify them as dense, moderate, or light. For each case, the histological stacks and Neurolucida chartings from serial coronal sections were manually aligned and reconstructed into 3D models using IMOD (Boulder Laboratory for 3D Electron Microscopy) ^41^. The outlines from all 3 cases were then aligned onto a common template developed in-house^40^ using an anatomical landmark-based registration in IMOD and stacked to produce a 3D model of the bundles. For each bundle, we generated centroids in template space. The white matter anatomical boundaries of the SLF-I and CB were outlined manually in template space following the definitions and visualizations in^8^.

### MRI acquisition

#### *Ex vivo* NHP data

The *ex vivo* macaque datasets include dMRI for 4 whole brains (2 *Macaca Mulatta, 2 Macaca Fascicularis*) acquired in a small-bore 4.7T Bruker BioSpin scanner at two different spatial resolutions. Specimens had been previously fixed in sucrose as described in ^38,42^ and were placed in a sealed Fomblin-filled plastic bag for scanning to eliminate air bubbles that would cause susceptibility artifacts. We collected 515 diffusion-weighted (DW) volumes (maximum b = 40,000 s/mm^2^) corresponding to a cubic lattice in q-space, including one b=0 volume. For dataset M3 a multi-slice multi-echo (MSME) was also acquired using the Carr Purcell Meiboom Gill (CPMG) sequence ^43^. Acquisition details are reported in Supplementary Table 3.

#### *Ex vivo* human data

The two *ex vivo* left hemispheres used in this study were extracted from two human brains (for demographic and clinical details, see supplementary table 4), which had been obtained from the Massachusetts General Hospital Autopsy Suite and fixed in 10% formalin for at least two months. Both subjects were neurologically normal prior to death; only mild hypoxic ischemic changes and mild hypertension were diagnosed for specimen 1 and 2 respectively.

The hemispheres were transferred into sealed plastic bags filled with liquid Fomblin and imaged in a 3T Siemens Trio scanner at 750 μm isotropic spatial resolution using a product 32-channel head coil and a 3D diffusion-weighted steady state free precession (DW-SSFP) sequence ^44^. We collected 60 DW volumes (effective b=3,373 s/mm^2^ ^45^) and 8 b=0 volumes (for acquisition details, see supplementary table 4). The frontal lobe of each hemisphere was then blocked into coronal slabs of approximately 1 cm thickness and a small block (roughly 2x1x2 cm) was excised from the dorsomedial section of each slab (Figure 9). These blocks included the white matter underlying the superior frontal gyrus, which was expected to contain SLF-I fibers. We obtained 3 blocks for specimen 1 and 4 blocks for specimen 2. Each of these samples was immersed in a Fomblin-filled syringe and all air was removed from the syringe to avoid susceptibility artifacts. Samples were scanned in a small-bore 9.4T Bruker Biospec system at either 250 μm (Specimen 1) or 300 μm (Specimen 2) isotropic spatial resolution. For all blocks, we collected 515 volumes, corresponding to points on a 11^3^ cubic lattice in space enclosed in a sphere, with b_max_ = 40,000 s/mm^2^ (supplementary table 4). All blocks were allowed to reach room temperature for at least 3 hours before scanning. After MRI acquisition, the blocks were prepared for PS-OCT as described in section 2.4.

**Figure 9.**
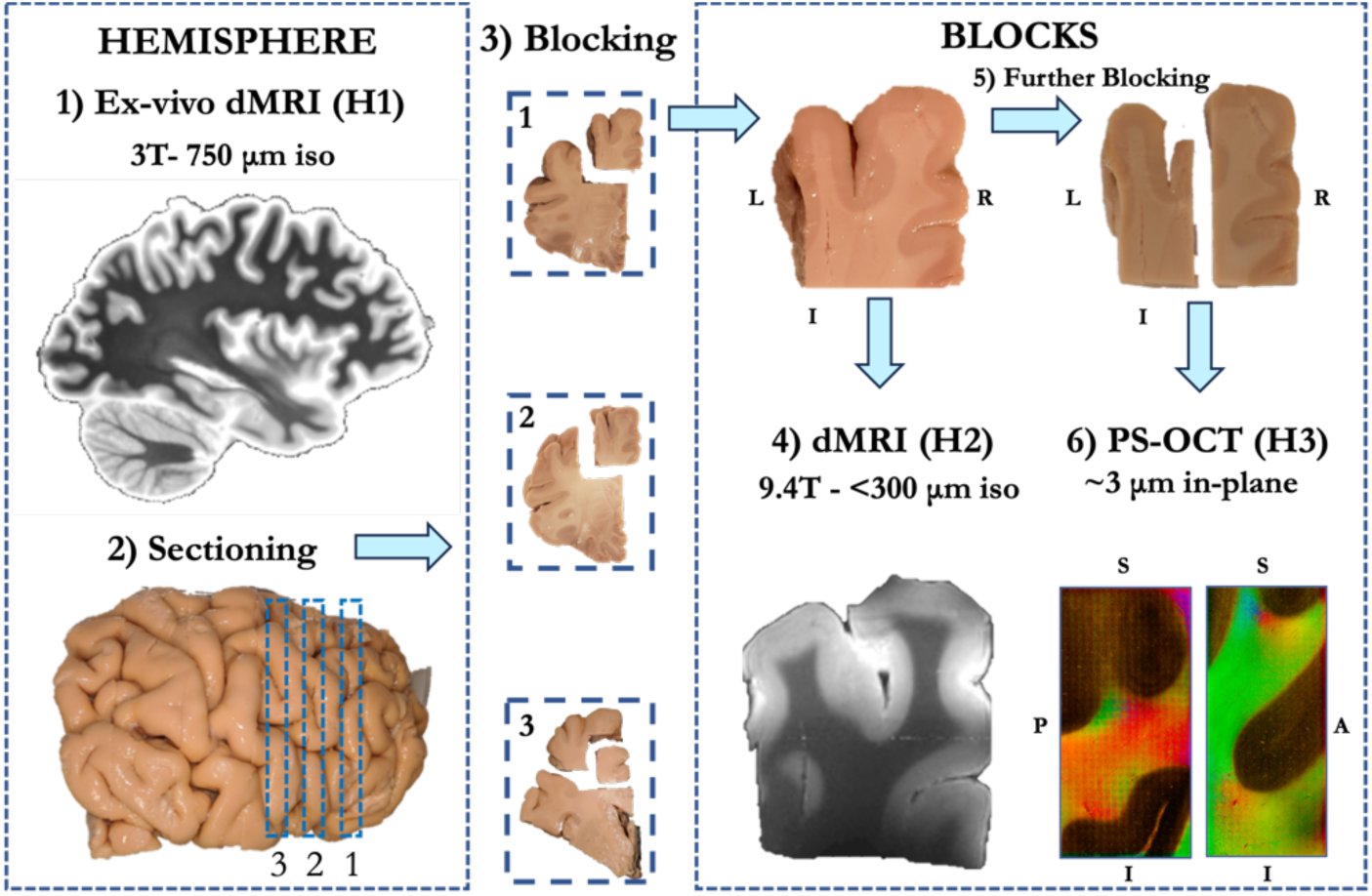
*Ex vivo* human datasets H1-H2-H3 workflow. The brain hemispheres are first imaged on a 3T Siemens Trio system at 750 μm (**1**) (Dataset H1). After scanning, each hemisphere is blocked into 1-cm-thick coronal slabs (**2**). Smaller blocks are then cut from the dorso-medial portion of each slab within the superior frontal gyrus (**3**) and scanned in a 9.4 T small-bore Bruker scanner at either 250 μm or 300 μm isotropic spatial resolution (**4**) (Dataset H2). Following this second scan, blocks are further reduced in size (**5**) for PS-OCT imaging (**6**) (Dataset H3).

#### *In vivo* human data

We used publicly available *in vivo* human datasets from eleven healthy subjects acquired on the 3T MGH Connectom 1.0 Scanner. (Acquisition details are reported in supplementary table 4.) Ten of these subjects were scanned as part of the MGH-USC Human Connectome Project (7 males; mean age: 31.4 s± 9.2 years; age range: 20 to 59 years) using 2D echo-planar imaging (EPI). Data were acquired at 1.5 mm spatial resolution, with 552 DW volumes distributed over four shells (64 b = 1000, 64 b = 3000; 128 b = 5000; 256 b = 10000 s/mm^2^) and 40 b=0 s/mm^2^ volumes^19^. One subject (male, 32 years) was scanned using a signal-to-noise ratio–efficient simultaneous multislab imaging technique (gSlider-SMS) with gSlider factor=5, across nine two-hour sessions ^46^. Data were collected at 760 μm spatial resolution, with 2664 DW volumes distributed over two shells (420 b=1000, 840 b=2500 s/mm^2^, along with paired volumes with reverse phase-encode direction) and 144 b=0 s/mm^2^ volumes^18^. For all subjects, a T1-weighted image was also acquired at 1 mm isotropic resolution (supplementary table 4).

### MRI Processing

#### *Ex vivo* macaque data

Data were preprocessed as previously described ^47^ using a pipeline that included MP-PCA denoising^48^ and correction for Gibbs ringing ^49^ in MRtrix3 ^50^, and correction for motion and eddy-current distortions in FSL ^51,52^. The diffusion tensor model was also fit to the volumes with b ≤ 4000 s/mm^2^using DTIFIT to obtain fractional anisotropy (FA) maps. For CSD reconstruction, we resampled data from a grid to shells in q-space using a fast implementation of the non-uniform fast Fourier transform (NUFFT) ^53^. We resampled the data onto two shells (b = 6,000 and 12,000 s/mm^2^) and fit fiber orientation distribution functions (fODFs) to the pre-processed data using a multi-shell, multi-tissue (MSMT)-CSD algorithm (l_max_ = 8) ^36^ with two tissue types (white and grey matter) in MRtrix3. The labels of the cortical parcellation were obtained using a manually edited, volumetric version of the Paxinos atlas ^54^ developed in-house on blockface microscopy images and aligned to the NMT v2.0 space. Briefly, anatomical boundaries were manually delineated on every fourth blockface section of an in-house common macaque template using iMod (Boulder Laboratory for 3D Electron Microscopy)^41^. To generate a complete volumetric representation of the atlas, interleaving sections were interpolated in three dimensions using the c3d toolkit from ITK-SNAP. Blockface images were then converted to NIfTI format and aligned to the asymmetric NMT v2.0 space in ANTs ^55^, by first registering the blockface volume to a structural MRI scan of the same animal, and then aligning this to the NMT v2.0 template. The resulting transformation matrices from the registration pipeline were concatenated and used to warp the annotated labels into NMT space. To map these labels to the dMRI space, the 0-th order spherical harmonic image from the CSD reconstruction of each macaque dataset was mapped to the NMT v2.0 template ^56^ by means of symmetric diffeomorphic registration in ANTs^55^. The inverse registration was then used to warp the anatomical labels from the NMT v2.0 space to the dMRI space of each macaque dataset. The same set of transformations were used to map the white matter masks of the SLF-I and CB from the in-house blockface template to the dMRI space of each macaque dataset. Surfaces and segmentations of white matter tissue were obtained using the FreeSurfer recon-all pipeline modified for NHP data ^57^. For dataset M2, the pipeline was run on the mean diffusion-weighted image (average of all volumes with b-value > 0), normalized by the b = 0 image. For dataset M3 it was applied to T2-weighted multi-slice multi-echo (MSME) images that had been collected in the same session as the dMRI data.

#### *Ex vivo* human data

Data from both hemispheres (dataset H1) and blocks (dataset H2) were denoised using MP-PCA in MRtrix3 ^48,50^ and corrected for eddy current-induced distortions in FSL ^52^. The diffusion tensor model was also fit to the data using DTIFIT to obtain FA maps (for dataset H2 we used only volumes with b-value ≤ 4000 s/mm^2^). For dataset H1, which consists of only one shell, fODFs were estimated in MRtrix3 using the *Tournier* CSD algorithm (l_max_ = 6) ^58^. The effective b-value (3,373 s/mm^2^) used in the estimation was computed as described in ^45^. All diffusion weighted volumes (b-value > 0) were averaged and normalized by the b-value = 0 volume to create an image that was processed in FreeSurfer v.7.5.0 ^59,60^ to obtain automated parcellations of the cortex through the *recon-all* stream ^61–64^. To maximize correspondence between human and NHP prefrontal areas, we used a publicly available parcellation scheme that translated the anatomical definitions of cytoarchitectonic regions from Petrides et al. 2012 ^21^ to the cortical surface of the *fsaverage* template ^22^. We mapped that parcellation from the *fsaverage* surface to the individual surface using the inverse of the FreeSurfer spherical morph^65^. No additional registration was needed as recon-all stream was run in individual diffusion space using the mean diffusion-weighted image (average of all volumes with b-value > 0), normalized by the b = 0 image. For CSD reconstruction on the dMRI data for the blocks (dataset H2), we resampled data from a grid to shells in q-space as previously described ^66^, using the same NUFFT implementation ^53^ as for the *ex vivo* macaque datasets M2, M3. We generated data on three shells over 256 diffusion directions (64 b=4,000; 64 b=8,000; 128 b=12,000 s/mm^2^), roughly corresponding to the following *in vivo* b-values: b=1000, 2000, 3000 s/mm^2^ ^67^. We estimated fODFs in MRtrix3 using the MSMT-CSD algorithm and tissue-specific response functions obtained manually from selected white and gray matter voxels (l_max_ = 8) ^50^.

#### *In vivo* human data

For all *in vivo* datasets we obtained preprocessed, publicly available data ^18,68^ and reconstructed fODFs in MRtrix3 using an MSMT-CSD algorithm (l_max_ = 8) ^37^. The diffusion tensor model was fit to the data from the b = 1000 s/mm^2^ shell using DTIFIT to obtain FA maps. The T1-weighted images from all subjects were processed in FreeSurfer v.7.5.0 to obtain automated cortical parcellations using the *recon-all* stream ^61–64^. To maximize correspondence between human and NHP prefrontal areas, we used a publicly available parcellation scheme that includes the cytoarchitectonic regions from Petrides et al. 2012 ^21^ drawn onto the *fsaverage* surface template ^22^. We mapped this parcellation from the *fsaverage* surface to the individual surface using the inverse of the FreeSurfer spherical morph, and then used a boundary-based, affine registration method ^69^ to align the dMRI b=0 image to the T1-weighted image and map the parcellation to the dMRI volume space using the inverse of that transform.

### PS-OCT acquisition and preprocessing

Following MRI scanning, the blocks obtained from each human hemisphere were imaged using a custom-built automatic serial sectioning PS-OCT system ^70^. The system integrated a commercial spectral domain PS-OCT centered at 1300 nm, motorized xyz translational stages, and a vibratome to section the tissue block. Automatic imaging and sectioning of brain blocks is controlled by custom-built software written in C++ for coordinating data acquisition, xyz stage translation, and vibratome sectioning. Blocks were embedded in agarose to protect against shear forces during tissue cutting, and then imaged in phosphate-buffered saline (PBS) immersion. Before embedding, each tissue block from specimen 1 was first further cut into two smaller blocks (Figure 9) to fit the capacity of the vibratome. Block 1 and 2, corresponding to the dorso-medial prefrontal cortex (dmPFC), were divided into a medial and lateral portion, while block 3 was divided into an anterior and posterior section. From specimen 2 we only divided block 3 into smaller anterior and posterior blocks, as an upgrade to the system increased the capacity of the vibratome allowing it to section and image bigger samples. The samples were flat faced with the vibratome and then imaged using an objective (OCT-LSM3, Thorlabs, Newton, NJ). The PS-OCT imaging/slicing plane for all blocks was sagittal, so that SLF fibers would be mainly in-plane. Serial scanning was performed using a snaked tile configuration scheme, with each tile having a FOV of 1.1 mm^2^ and 550 μm overlap between adjacent tiles for blocks from specimen 1, and a FOV of 3 to 4 mm^2^ and 500 μm overlap between adjacent tiles for blocks from specimen 2. The slice thickness was 100 μm for all samples. Specific acquisition parameters for each block are provided in supplementary table 4, along with sample size and in-plane resolution. Using a Jones matrix formalism, the retardance and the optic axis orientation were computed from the amplitude and the phase of the complex signals, respectively, on the two polarization channels ^71^. For each of these two modalities, data were saved as 2D en-face maps and image tiles were stitched using the Fiji software ^72,73^. The enface slices were then stacked to form a volume. As the tissue blockface is imaged before it is cut, the images are undistorted and can be stacked across sections without further inter-slice registration. For visualization of en-face optic axis orientation slices, HSV images were created using the orientation measurement to determine the hue/color in each pixel, and the retardance to modulate color intensity. Colored orientation images were despeckled using Fiji ^72^. The PS-OCT data from blocks acquired at a resolution higher than 10 μm were all downsampled to 10 μm in-plane resolution to facilitate further analyses. The optic-axis orientation vectors obtained from PS-OCT represent axonal orientations within the sagittal imaging plane, and can be directly used to perform tractography as explained in section 2.7.1.

### Registration of dMRI and PS-OCT human data

The PS-OCT image stacks (dataset H3) were aligned to the dMRI scan from the same tissue block (dataset H2), which was aligned to the dMRI scan of the respective hemisphere (dataset H3). We performed the PS-OCT to dMRI alignment by registering the retardance map from the former to the FA map from the latter, using a robust affine registration framework ^74^. The use of affine registration is justified by the absence of non-linear distortions in PS-OCT compared to other histological techniques, and was previously assessed by our group^75^. We aligned each block to the respective hemisphere manually using both the FA maps and the b = 0 s/mm^2^ volumes from the respective dMRI datasets. We then concatenated the transformations to map the PS-OCT blocks to their respective hemisphere.

### Common parcellation scheme

We created a common parcellation scheme for macaque and human MRI data to ensure homology (Supplementary Table 1). For the macaque data, dorsomedial labels were manually edited from a volumetric version of the Paxinos atlas^54^ developed in-house, as described in *MRI processing, ex vivo macaque data*. For the human data, we combined the FreeSurfer parcellation with a publicly available parcellation scheme that translates anatomical definitions of cytoarchitectonic regions of the prefrontal cortex to the *fsaverage* cortical surface as described in the *MRI processing* sections. For each area in the human parcellation we identified the corresponding area(s) in the macaque parcellation (Supplementary Table 6).

### Tractography analyses

We performed qualitative and quantitative analyses to assess the connectional anatomy of the SLF-I across species and scales. For qualitative analyses, we performed manual virtual dissections using deterministic tractography across scales in humans and probabilistic tractography in macaques. For quantitative analyses, we computed connectivity matrices between dorso-medial cortical regions using probabilistic tractography in macaques and humans, and generated connectograms to visualize them.

#### Manual virtual dissection of the SLF-I

Virtual dissections of SLF-I connections from high-resolution *ex vivo* data (H1, H2, H3, M2, M3) were performed manually in Trackvis by author C.M. To integrate the information across the different dMRI and PS-OCT human datasets (H1, H2, H3), we developed a multi-scale tractography algorithm that can use orientation datasets that come from the same brain but that are available at different resolutions and within different FOVs. This framework includes a tractography algorithm specifically developed in-house for microscopic data that we applied to the PS-OCT orientation maps in this study, with a search angle of 10° and a search radius of 15 voxels ^76^. Streamlines were generated in the native space of each dMRI or PS-OCT dataset. For the human ex vivo datasets, these were then mapped to the respective whole-hemisphere dMRI space (Dataset H1), where all virtual dissections were carried out. For macaque data, one sample from M2 (700 μm) and one from M3 (500 μm) were selected for virtual dissection. Whole-brain local probabilistic tractography was performed in MRtrix3 (angle threshold = 45°; step-size = 0.5 x voxel size; maximum length = 100 mm; cutoff = 0.1) by randomly placing 10 seeds per voxel within a whole-brain white matter mask. Virtual dissections were carried out by manually drawing a series of coronal inclusion ROIs along the dorso-medial white matter of the brain to identify all the streamlines running within the full extent of the SLF-I. The anatomical definition of the SLF-I from ^8^ was used to define the borders of the ROIs. Exclusion ROIs were placed to reject streamlines projecting to brain regions outside the dorso-medial cortex. In addition to the SLF-I streamlines, we isolated streamlines belonging to neighboring white matter bundles including the CB and its projections, the corona radiata (CR), and neighboring fibers connecting to adjacent lateral regions.

#### Connectivity matrices of SLF-I and neighboring fibers

We generated connectivity matrices from each whole-brain or hemisphere dMRI dataset included in this analysis (H1, H4, H5, M2, M3). Local probabilistic anatomically constrained tractography (ACT) was performed in MRtrix3 ^77^ by randomly seeding within each voxel in a white matter mask (angle threshold = 45°; step-size = 0.5 x voxel size; maximum length = 150 mm for humas, 100 mm for NHP; cutoff = 0.1). The number of seeds per voxel was 10 for H1 and H4, and 50 for M2, M3, and H5, to ensure a comparable number of streamlines per unit volume across different spatial resolutions and brain sizes. The white matter mask used for seeding included the dorso-medial frontal and parietal white matter (including SLF-I fibers) and the cingulate white matter (including CB fibers). This mask was obtained by extracting and aggregating labels from the white-matter parcellation (*wmparc*) that is produced in FreeSurfer recon-all by labeling white matter voxels with the closest cortical label (max distance = 5 mm). For the SLF-I, the following white matter labels were extracted: paracentral, precentral, precuneus, superior frontal, superior parietal. For the CB, the following labels were extracted: caudal anterior cingulate, rostral anterior cingulate, posterior cingulate, isthmus cingulate. The anatomical accuracy of these masks was visually inspected for each sample and masks were manually edited in freeview if needed. For all included datasets, tractography was constrained to a binary mask including the dorsomedial cortical regions of interest listed in Table 7 along with the underlying white matter. As our goal was to analyze association fibers, we excluded superficial white matter fibers, which would confound this analysis, using criteria based on length and distance from the cortex. Specifically, we discarded all streamlines shorter than 30 mm in humans ^78^ and 10 mm in NHP. We also created a mask of the superficial white matter in FreeSurfer by sampling between the white matter surface and 2 mm into the white matter in humans and 0.5 mm in NHP. We subtracted this superficial white matter volume from the whole-brain white matter segmentation obtained from the recon-all stream. Only streamlines traversing this deep white matter mask were included in the final tractograms. For each dataset, two sets of tractograms were generated: one including streamlines coursing through the CB and one excluding streamlines coursing through the CB. For each tractogram, connectivity matrices were then generated in MRtrix3 ^50^ by projecting streamlines radially by 2 mm from the streamline endpoint in search of a parcellation node voxel.

## Author contributions

C.M. contributed to conceptualization, methodology, MRI data acquisition, validation, formal analysis, investigation, manual annotations of dMRI and tracer data, and wrote the original draft, with inputs from all authors. S.H. and J.L. contributed to conceptualization, tracer data acquisition, histology data processing and manual annotations. M.R.C. contributed to investigation and analysis. A.Y. contributed to conceptualization, methodology, acquisition, and project supervision. All authors contributed to manuscript revision and approved the submitted version.

