## Supplementary Information for "The modular nature of long-association pathways in the human brain"

**Supplementary Tables**

| ***NHP*** | ***Human*** | | ***Anatomical Definition*** |
| --- | --- | --- | --- |
| ***Paxinos*** | ***FS Parcellation*** | ***Petrides*** |  |
| Area 10M |  | Area 10M | Anterior Prefrontal Cortex, Frontal Pole |
| Area 9M |  | Area 9M | Dorsomedial Prefrontal Cortex |
| Area 9L |  | Area 9L | Dorsolateral Prefrontal Cortex |
| Area 9/46d |  | Area 9/46d | Dorsolateral Prefrontal Cortex |
| Area 8B |  | Area 8B | Dorsomedial Prefrontal Cortex |
| Areas 6M (SMA), F2, F7 |  | Area 6 | Premotor |
| Area 4 |  | Area 4 | Primary Motor Cortex |
| Areas 24a, 24b, 24c, 24d, 32, 8/32, 9/32, 6/32 |  | Area 24, 32 | Anterior Cingulate Cortex |
| Areas 1, 2 (SS), 3a, 3b | Postcentral, Paracentral |  | Posterior Central Gyrus |
| Areas 23a, 23b, 23c, 31 | Posterior Cingulate, Isthmus Cingulate |  | Posterior Cingulate Cortex |
| PGM/31 | Precuneus |  | Precuneus |
| PE, PE Caudal, PE Cing., PEa | Superiorparietal |  | Superior Parietal Lobule |

**Supplementary Table 1. Common parcellation scheme of the dorsomedial cortex in macaque and human dMRI datasets.** For the macaque brains, we used a volumetric version of the Paxinos atlas (1) in NMT v2.0 template space (2) developed in-house and manually edited. For the human brains, we matched these parcels to a cortical parcellation that combines the frontal parcels of the Petrides parcellations scheme (3) with the posterior parcels from the *recon-all* stream. The table also reports the corresponding architectonic region in humans for each area.

| ***Anatomic tracer injections – Dataset M3*** | | | | | | | |
| --- | --- | --- | --- | --- | --- | --- | --- |
| ***Case #*** | **1** | **2** | **3** | **4** | **5** | **6** | **7** |
| ***Injection Site*** | dmPFC | dmPFC | FEF | Premotor | Premotor | Premotor | Parietal |
| ***Area*** | 9M | 9M | 8B | 6M | 6M | F7 | PE |
| ***Species*** | MN | MN | MR | MR | MR | MF | MR |
| ***Tracer*** | LY | LY | LY | FS | FS | FR | FR |

**Supplementary Table 2. Anatomic tracer injection cases.** For each of the 7 cases, the table reports the anatomical region and specific Brodmann area of the injection site, along with the species of the animal (MN, MR, or MF) and the type of injected tracer. dmPFC: dorsomedial prefrontal cortex; FEF: frontal eye field; FR: fluororuby; FS: fluorescein; LY: lucifer yellow; MF: Macaca Fascicularis; MN: Macaca Nemestrina; MR: Macaca Rhesus.

| **Specimen #** | **Hemisphere** | **Age (years)** | **Sex** | **PMI (hrs)** | **Cause of death** |
| --- | --- | --- | --- | --- | --- |
| **1** | LH | 70 | M | 24 | Coronary artery disease |
| **2** | LH | 60 | F | 2 | Adenocarcinoma of pancreas |

**Supplementary Table 3. Demographics for human brain specimens.** LH = left hemisphere; PMI = post-mortem interval.

**MRI Acquisition Parameters Tables:**

| ***MRI Acquisition Parameters NHP Datasets*** | | | |
| --- | --- | --- | --- |
| ***Dataset #*** | **M2** | **M3** | **M3** |
| ***Field Strength*** | 4.7 T | | |
| ***Sequence*** | 3D EPI | | MSME |
| ***Spatial resolution (mm^3^)*** | 0.7 | 0.5 | 0.5 |
| ***TR (ms)*** | 750 | 500 | 3000 |
| ***TE (ms)*** | 43 | 48 | 8: 8: 160 |
| ***G_max_ (mT/m)*** | 660 | |  |
| ***b-values (s/mm^2^)*** | b = 40,000 (b_max_) | | N/A |
| ***Gradient pulse width (ms)*** | 15 | 11 | N/A |
| ***Diffusion Time (ms)*** | 19 | 21 | N/A |
| ***FOV (mm)*** | 72x72 | 58x48 | 58x48 |
| ***PE*** | LR | LR | LR |
| ***Scan Time (hrs)*** | ~48 | ~47 | ~10 |

**Supplementary Table 4. dMRI acquisition parameters for NHP datasets M1 and M2.** Abbreviations: EPI = echo planar imaging; GRAPPA = generalized autocalibrating partial parallel acquisition; δ = gradient pulse width; ∆ = diffusion time; beff = effective b-value; bmax = maximum b-value; FOV = field of view; Gmax = maximum gradient strength; MSME = multi-sclice multi-echo; PE = phase encoding; SRmax= maximum slew rate; T1w = T1 weighted; TE = echo time; TR = repetition time.

| ***MRI Acquisition Parameters Human Datasets*** | | | | | | |
| --- | --- | --- | --- | --- | --- | --- |
|  | ***In Vivo*** | | | | ***Ex Vivo*** | |
| ***Dataset #*** | **H4** | | **H5** | | **H1** | **H2** |
| ***Field Strength*** | 3 T | | 3 T | | 3 T | 9.4 T |
| ***G_max_ (mT/m)*** | 300 | | 180 | |  | 480 |
| ***Contrast*** | Diffusion | T1w | Diffusion | T1w | Diffusion | Diffusion |
| ***Sequence*** | 2D EPI | MEMPRAGE | 2D EPI | MEMPRAGE | 3D DW-SSPF | 3D EPI |
| ***Resolution (mm^3^)*** | 1.5x1.5x1.5 | 1x1x1 | 0.76x0.76x0.76 | 1x1x1 | 0.75x0.75x0.75 | 0.25x0.25x0.25-0.34x0.34x0.34 |
| ***TR (ms)*** | 8800 | 2530 | 3500 | 2350 | 30.21 | 750 |
| ***TE (ms)*** | 57 | 1.15 | 75 | 1.61 - 7.19 | 25.12 | 50-72 |
| ***b-values (s/mm^2^)*** | 1000, 3000, 5000, 10000 | N/A | 1000, 2500 | N/A | 3,373 (b_eff_) | 40000 (b_max_) |
| ***δ (ms)*** | 12.9 | N/A | 6.4 | N/A | N/A | 15 |
| ***∆ (ms)*** | 21.8 | N/A | 30.8 | N/A | N/A | 21-30 |
| ***FOV (mm)*** | 210x210 | 256x256 | 290x288 | 256x256 | 203x216 | 34x34 |
| ***GRAPPA*** | 3 | 2 | 3 | 2 | 1 | 2 |
| ***MB*** | 1 | 1 | 2 | 1 | 1 | 1 |
| ***PE*** | AP | AP | AP | AP | RL | RL |
| ***Scan Time (hrs)*** | 1.5 | 0.1 | ~15 | 0.1 | ~30 | ~48 |

**Supplementary Table 5. MRI acquisition parameters for human datasets.** Abbreviations: EPI = echo planar imaging; GRAPPA = generalized autocalibrating partial parallel acquisition; δ = gradient pulse width; ∆ = diffusion time; b_eff_ = effective b-value; FOV = field of view; G_max_ = maximum gradient strength; MB = multiband; MEMPRAGE = Multi-echo Magnetization-Prepared Rapid Acquisition Gradient Echo; PE = phase encoding; SR_max_= maximum slew rate; T1w = T1 weighted; TE = echo time; TR = repetition time.

| **PS-OCT Acquisition Parameters** | | | | | | |
| --- | --- | --- | --- | --- | --- | --- |
| **Specimen** | **Block #** | **Tiles/Slice** | **Slices/Sample** | **Sample size (cm)** | **In-plane resolution (μm)** | **Scan time (hrs)** |
| **1** | *B2-l* | 600 | 56 | 0.5x0.8x2 | 2.75 | 96 |
|  | *B2-m* | 665 | 58 | 0.5x1x2 | 3.67 | 68 |
|  | *B3-l* | 858 | 51 | 0.5x1.3x2 | 3.93 | 69 |
|  | *B3-m* | 765 | 55 | 0.5x1x2.5 | 3.93 | 81 |
|  | *B4-a* | 480 | 111 | 1x0.8x2 | 3.93 | 80 |
|  | *B4-p* | 722 | 87 | 0.9x0.9x2 | 3.93 | 85 |
| **2*** | *B2* | 207 | 325 | 3x1.5x5 | 10 | 90 |
|  | *B3a* | 456 | 92 | 0.4x0.7x2 | 3.93 | 70 |
|  | *B3p* | 24 | 185 | 2x0.9x2 | 10 | 25 |
|  | *B4* | 66 | 189 | 2x1.5x3 | 10 | 44 |
|  | *B5* | 63 | 171 | 1.5x1.5x2.5 | 10 | 43 |

**Supplementary Table 6. PS-OCT Acquisition parameters.** For each specimen, the table reports the block number, the number of imaged tiles per slice, the number of imaged slices per sample, the depth, the approximate sample size, the estimated lateral resolution, and the total scan time. *****Most of the blocks from specimen 2 were acquired on an upgraded version of the same as-PS-OCT system that allowed sectioning and imaging of bigger samples.

**Supplementary Figures**

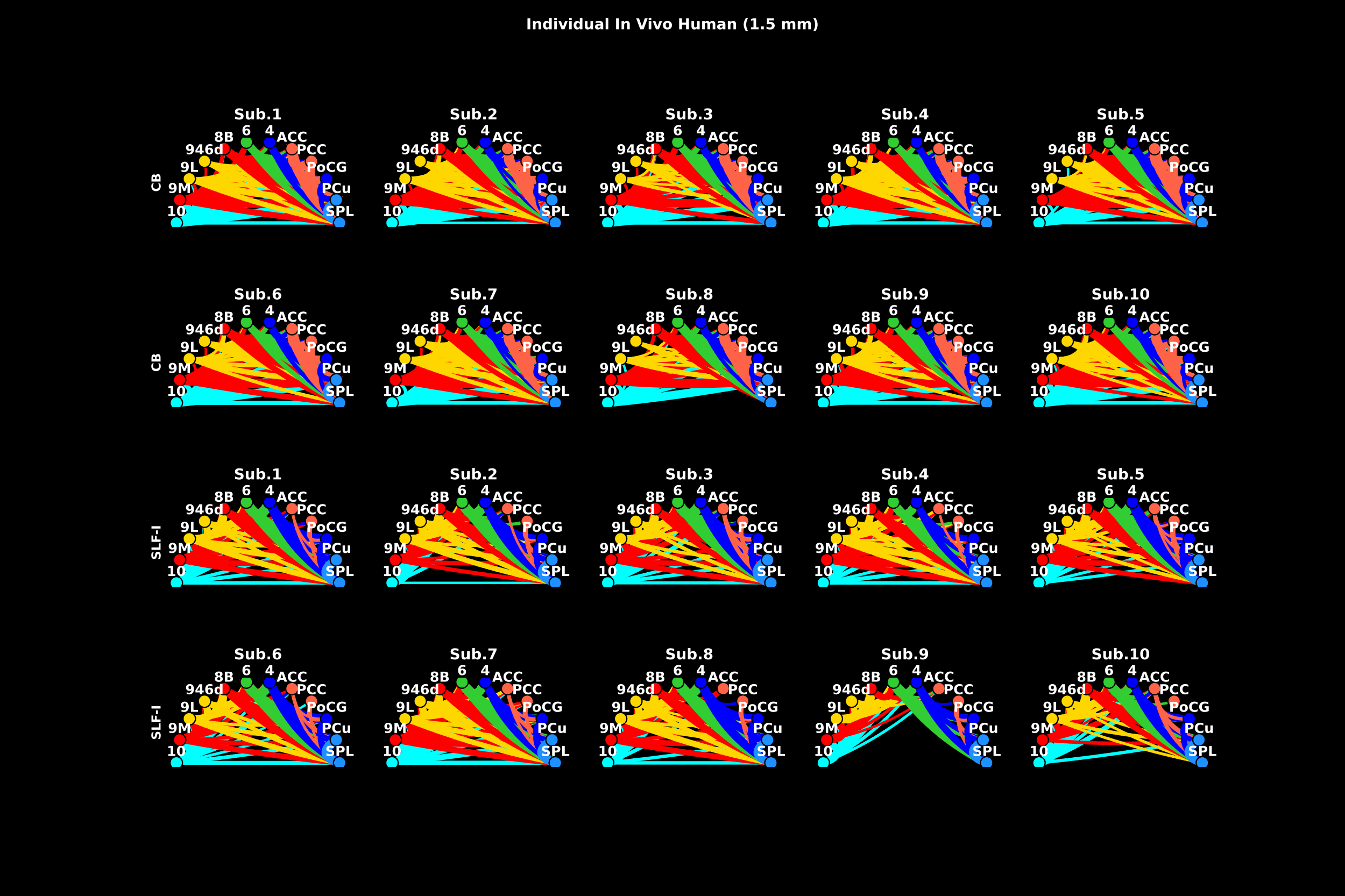

**Supplementary Figure 1.** Individual connectograms are shown for each of the ten subjects from dataset H5. The top two rows display connectivity profiles between dorsomedial cortical regions computed from streamlines coursing through the cingulum bundle, while the bottom row shows connectivity profiles computed from streamlines passing through the dorsomedial white matter where the SLF-I runs. The width of the lines is weighted by the number of streamlines between regions, and connection colors correspond to the cortical regions they link, arranged in anterior-to-posterior order: medial prefrontal cortex (mPFC, red) includes regions 9M and 8B; lateral PFC (lPFC, yellow) includes regions 9L and 9/46d; pre-supplementary motor area (pre-SMA, light green) corresponds to region 6; motor areas (blue) include region 4 and regions 1,2,3 in the posterior central gyrus (PCG) ; cingulate regions (orange) include anterior (ACC) and posterior cingulate cortex (PCC) (Areas 24, 32, 8/32, 9/32, 6/32, 23, 31); parietal region (purple) include the superior parietal lobule (SPL) and the precuneus (PCu).

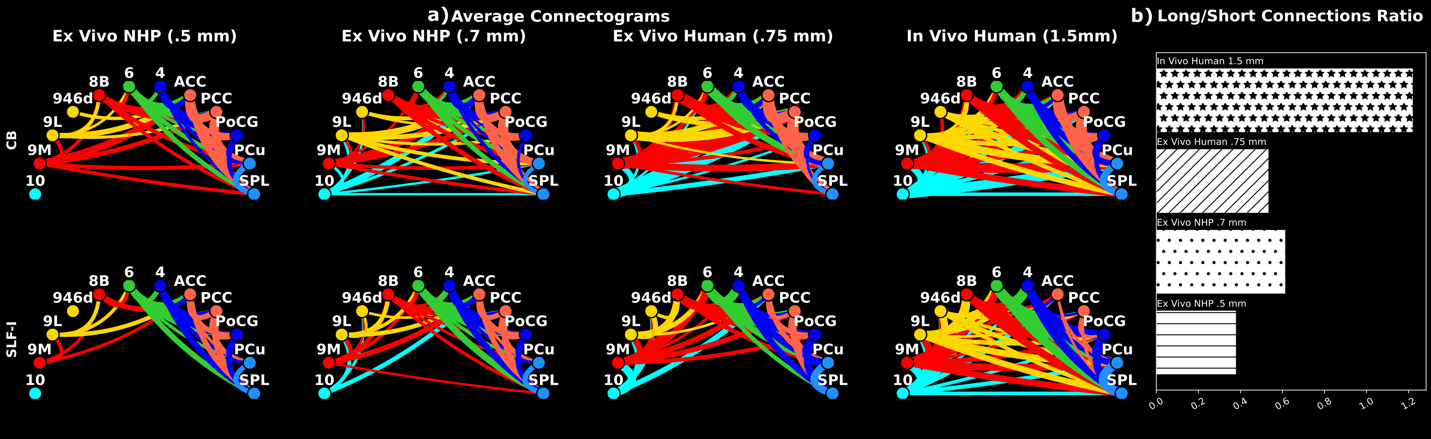

**Supplementary Figure 2. a)** Averaged connectograms are shown for each species and resolution. The top row displays connectivity profiles between dorsomedial cortical regions computed from streamlines coursing through the cingulum bundle, while the bottom row shows connectivity profiles computed from streamlines passing through the dorsomedial white matter where the SLF-I runs. The width of the lines is weighted by the number of streamlines between regions, and connection colors correspond to the cortical regions they link, arranged in anterior-to-posterior order: medial prefrontal cortex (mPFC, red) includes regions 9M and 8B; lateral PFC (lPFC, yellow) includes regions 9L and 9/46d; pre-supplementary motor area (pre-SMA, light green) corresponds to region 6; motor areas (blue) include region 4 and regions 1,2,3 in the posterior central gyrus (PCG) ; cingulate regions (orange) include anterior (ACC) and posterior cingulate cortex (PCC) (Areas 24, 32, 8/32, 9/32, 6/32, 23, 31); parietal region (purple) include the superior parietal lobule (SPL) and the precuneus (PCu). **b)** The ratio of long to short connections is shown for each of the averaged connectograms in a. A threshold equal to 3 nodes was used to separate short versus long connections within the connectogram.
